# Deep multiplexed 50-marker imaging of circulating tumor cells expands actionable biomarker profiling for precision oncology

**DOI:** 10.1101/2025.10.23.684075

**Authors:** Timothy J. Mann, Ye Zheng, Tanzila Khan, Tim Huang, Udit Nindra, Yafeng Ma, Alexander James, Daniel P. Neumann, Felix V. Kohane, Ihuan Gunawan, Fatemeh Vafaee, Wei Chua, Paul de Souza, Tara L. Roberts, Therese M. Becker, John G. Lock

## Abstract

Liquid biopsy-derived circulating tumor cells (CTCs) offer a minimally invasive avenue for precision oncology by enabling longitudinal monitoring of actionable molecular and cell-phenotypic biomarkers. However, conventional immunofluorescence imaging-based CTC profiling captures just 4-5 markers per cell, restricting crucial insights into oncogenic and resistance drivers, as well as tumor cell heterogeneity. Here we present an integrated pipeline employing deep multiplexed imaging to profile up to 50 molecular markers per CTC, capturing expression, phosphorylation, and subcellular localization of diverse biomarkers. Validated in a multi-stage prostate cancer resistance model and applied to prostate cancer patient-derived CTCs, this approach enhances CTC classification and reveals inter– and intra-patient heterogeneity correlating with therapy response. Machine learning identified therapeutically actionable signatures, suggesting patient-specific treatment strategies. This deep multiplexed imaging pipeline advances the utility of CTCs in guiding personalized cancer therapy by providing comprehensive molecular and phenotypic insights through minimally invasive liquid biopsies.

**Highlights:**

– Multiplexed immunofluorescence imaging of circulating tumor cells captures up to 50 molecular markers per cell with subcellular localisation.
– Single-cell and subcellular analyses identify quantitative signatures correlating with patient outcome, as well as therapeutically actionable biomarkers.
– This method transforms the utility of CTCs for future use in patient stratification and for guiding personalised therapies.

Graphical Abstract

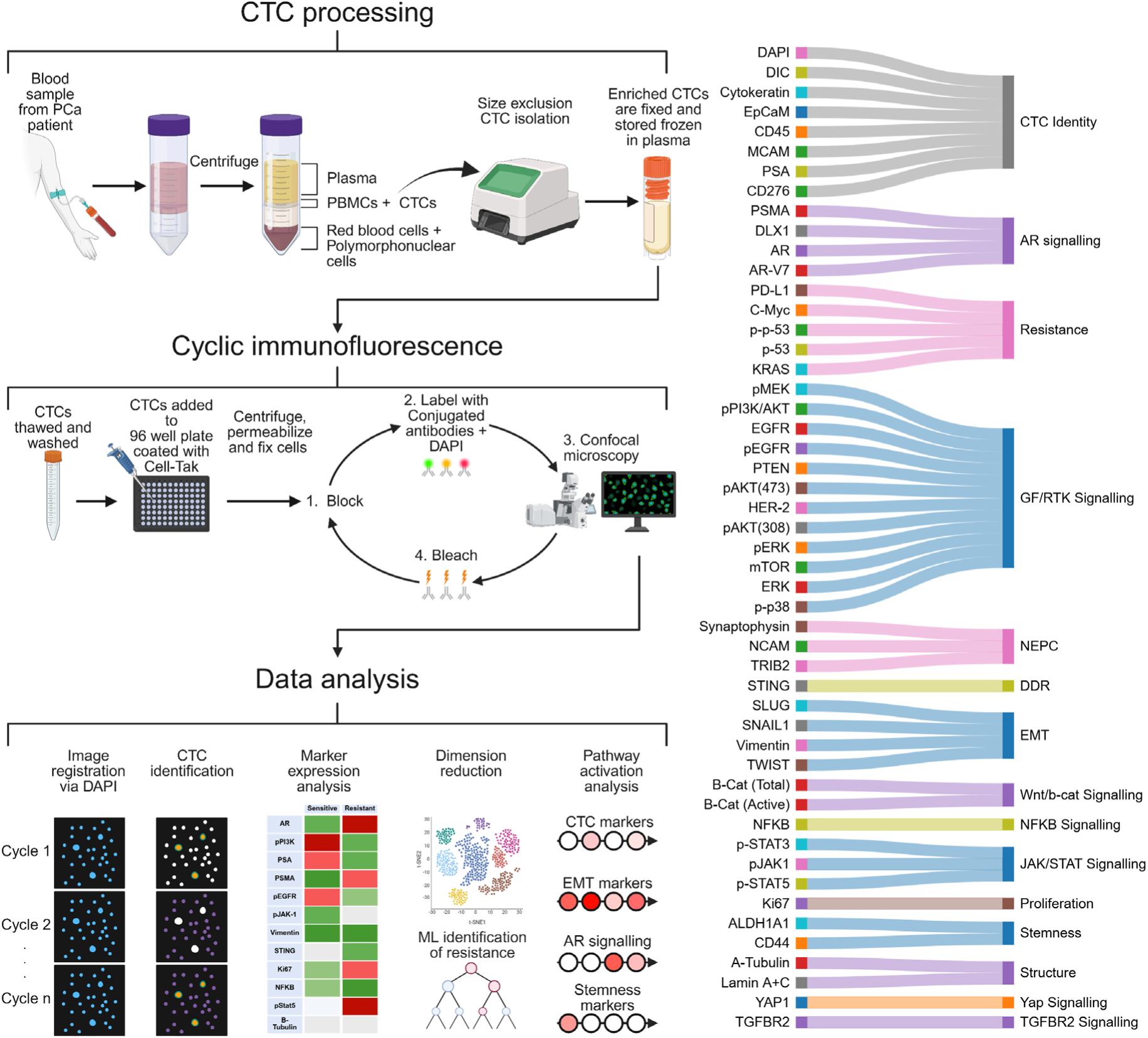

## Introduction

Precision oncology involves the molecular profiling of cancers to guide optimal therapies, increasing treatment efficacy while minimizing side-effects and futile treatment^1,2,3^. As the arsenal of targeted therapies expands, continually matching the right treatment to the right patient at the right time during cancer progression becomes an increasingly complex challenge. Yet such precision patient-to-therapy matching is vital to personalize and adapt patient management in clinical practice, as well as to enhance the success of clinical trials for targeted therapies impacting only a subset of patients^4^. Liquid biopsy-based blood sampling permits the analysis of circulating tumor cells (CTCs), providing a minimally invasive method for molecular-to-cellular scale cancer biomarker profiling that is, unlike solid biopsy analysis, safely compatible with extensive longitudinal sampling to track cancer adaptation and evolution^5^. CTCs play explicit roles in metastasis and CTC enumeration is currently emerging as a prognostic biomarker to stage patients with various cancers^6^. CTCs can also illuminate tumor heterogeneity by directly informing upon oncogenic signalling and cell state drivers at the functional, protein level^7^. For example, CTC-expression of key markers such as nuclear androgen receptor-variant 7 (AR-V7)^8^, programmed death-ligand 1 (PD-L1)^9^, prostate specific membrane antigen (PSMA)^10^, human epidermal growth factor 2 (HER-2)^11–12^ and estrogen receptor (ER)^13^ have predicted response to treatment in clinical trials^5^. Such molecular interrogation of CTCs is most commonly mediated through immunofluorescence (IF) imaging^5,14,15^. However, standard CTC IF imaging achieves detection of just 4-5 molecular markers per cell^16^. This limitation has shaped widespread adoption of a correspondingly constrained 3 or 4-plex ‘canonical definition’ of CTC identity: DAPI or Hoechst (nuclear)-positive; Cytokeratin and/or EpCAM (epithelial)-positive; CD45 (immune cell marker)-negative^17,18^. It is increasingly apparent that this ‘low-plex’ definition is insufficient to capture the true heterogeneity of CTCs^19–22^, undermining accurate enumeration of CTCs with non-canonical characteristics. At the same time, this 3-4 plex CTC definition in the context of standard 4-5 plex IF imaging leaves at most 1-2 additional molecular marker ‘channels’ available to further characterise CTCs in terms of their phenotypic diversity (e.g. in the epithelial-mesenchymal spectrum^23–26^) or to detect oncogenic signalling and/or resistance pathway biomarkers that could guide targeted therapy selection^27–31^. Yet precision oncology in heterogeneous cancers calls for far more extensive molecular profiling to simultaneously monitor numerous alternate phenotypic, oncogenic– and resistance-biomarkers to guide optimal selection of personalised therapies. The low-plexity of standard IF-based CTC analyses thus constitutes a pervasive and long-standing barrier to full exploitation of CTCs as sources of diverse molecular-to-cellular scale biomarkers that could guide precision oncology.

To address this widespread limitation, we have performed integrated development of an end-to-end methodological pipeline (including: *CTC enrichment; storage & retrieval; CTC surface attachment; cyclic labelling/imaging; image processing; data parsing; batch-normalized quantitative data analysis; machine learning for CTC classification & feature selection*) that enables deep multiplexed IF image profiling of CTCs using cyclic immunofluorescence (CycIF)^32^. We thus detect as many as 50 molecular markers per cell; ten-fold greater than standard CTC IF imaging methods in widespread use^33,34^. This enhances CTC identification and permits concurrent evaluation of numerous functional indicators (protein/phospho-protein levels and subcellular localizations) across multiple molecular pathways driving cancer progression and therapy resistance, while also delineating key cancer cell phenotypes (epithelial, mesenchymal, stem-like, proliferative). Such detailed molecular-to-cellular scale phenotyping of CTCs is especially vital in late-stage disease since tumor evolution and adaptation may render archival tissue samples (from early-stage solid biopsies) ‘out-of-date’ as sources of molecular guidance for further treatments^5^. Late-stage prostate cancer (PCa) exemplifies this challenge, as early PCa overwhelmingly relies on androgen receptor (AR) signalling and is thus sensitive to androgen deprivation therapy (ADT)^35^, however eventual ADT-resistance (castrate resistant prostate cancer, CRPC) can be driven by a variety of alternate oncogenic signalling and resistance pathways^36,37^. Above the molecular scale, cell phenotype changes such as neuroendocrine differentiation are also crucial mediators of therapy resistance^38^. Broad-spectrum profiling is thus vital in heterogenous late-stage PCa to parse in each CRPC patient which molecular driver/resistance pathways are active and which cancer cell phenotypes are present, so as to guide optimal therapy selection.

Accordingly, we here provide proof-of-principle for the utility of deep multiplexed CTC image profiling in late-stage PCa, with a view to enhancing capacity for minimally invasive guidance of precision oncology. We initially validated this approach using an established three-stage cell line model of acquired PCa therapy-resistance^37^, emulating longitudinal monitoring. We detected rich quantitative molecular signatures per cell including the levels and subcellular localizations of numerous biomarkers with known links to resistance. We next translated this approach to multiplexed imaging of size-enriched CTCs from PCa patient blood samples, finding molecular– and cellular-scale signatures correlating with patient clinical presentation. This unprecedented data revealed significant CTC heterogeneity between as well as within individual patients. Notably, intra-patient heterogeneity revealed reduced CTC population entropy (i.e. lower diversity) in therapy-resistant patients, suggesting the selection of specific resistance signals and/or phenotypes in these patients. In some patients, we observed multiple distinctive CTC subpopulations with different actionable targets; illuminating the future potential to guide combination treatment strategies to suppress distinct tumor cell subpopulations.

These proof-of-principle results suggest that deep multiplexed image profiling can enhance the identification of clinically actionable biomarkers in CTCs and may inform future studies aimed at guiding selection of current PCa and cancer-agnostic therapies. This novel strategy may also provide additional information relevant to patient stratification for emerging clinical trials. Overall, this approach appears poised to facilitate both fundamental and translational advances in understanding of the molecular and cellular mechanisms driving cancer progression, therapy resistance and metastasis; thereby advancing the emerging paradigm of precision oncology.

## Methods

### Sample collection

All reagents, their sources and identifiers are listed in Table 1.

**Table 1:** Reagents, Sources and Identifiers.

| REAGENT or RESOURCE | SOURCE | IDENTIFIER |
| --- | --- | --- |
| DMEM | Gibco | 11995-065 |
| L-Glutamine | Merck | 59202C-100ML |
| Penicillin/Streptomycin | Sigma-Aldrich | P4333 |
| Trypsin/EDTA | Sigma-Aldrich | T2601 |
| EDTA vacutubes | Greiner Bio-One | #455036 |
| Enzalutamide | Sapphire Biosciences | MDV3100 |
| PFA | Electron Microscopy Services | 15713 |
| DMSO | Sigma-Aldrich | D2650-100ML |
| CoolCell LX | Merck | CLS432001 |
| 96 well glass bottom plate | Cellvis | #:P96-1.5H-N |
| Cell Tak | Corning | 354240 |
| NaHCO<sub>3</sub> | Chem-Supply | SA-001 500g |
| Cryovials | Rowe Scientific | PC0345 |
| Triton X-100 | Thermofisher | HFH10 |
| BSA | Sigma-Aldrich | A7906-500g |
| DAPI | Thermofisher | D1306 |
| ibiseal cover film | Ibidi | 10874 |
| NaOH | Sigma-Aldrich | S8045 |
| H<sub>2</sub>O<sub>2</sub> | Univar | AJA260-500ML |
| Lymphoprep | Stemcell Technologies | #07851/07861 |
| Sepmate-50 | Stemcell Technologies | 85450 |
| BioTek 50TS Microplate washer | Millipore | 40-301 |
| Genesis Single-Cell System | Biorad | 12019822 |
| LNCaP | Gifted | CVCL_1379 |
| DU145 | Gifted | CVCL_C0QR |
| V16D | Gifted | Kind gift from Prof. Amina Zoubeidi |
| MR49F | Gifted | Kind gift from Prof. Amina Zoubeidi |
| Mycoalert plus mycoplasma kit | Lonza | LT07-710 |
| NIS Elements | Nikon | https://www.microscope.healthcare.nikon.com/products/software/nis-elements/software-resources |
| FIJI 1.54 | NIH | https://imagej.net/software/fiji/downloads |
| Cell profiler 4.2.6 | Broad Institute | https://cellprofiler.org/previous-releases |
| KNIME 5.2 | KNIME | https://www.knime.com/downloads |
| Python 3.12 | Python Software Foundation | https://www.python.org/downloads/release/python-3120/ |
| R 4.3.1 | The R Foundation | https://cran.r-project.org/bin/windows/base/old/ |

#### Patient Sample collection

Seven patients (age: 69-88) with advanced, metastatic PCa, and expected high CTC counts were recruited from Liverpool hospital and St. George private hospital, Sydney, Australia. Written informed consent was obtained from all patients under the South Western Sydney Local Health District Human Research Ethics Committee approved protocol HREC/13/LPOOL/158. Single blood samples were collected in 9 mL EDTA vacutubes.

#### Cell lines

Prostate cancer cell lines LNCaP, DU145 were STR-authenticated (ARGF, Melbourne, Australia) and tested for mycoplasma (MycoAlert^TM^ PLUS Mycoplasma Detection Kit, Lonza) and LNCaP –derived V16D and MR49F^37^ (a gift from Dr. Amina Zoubeidi, Vancouver Prostate Centre, Canada) were cultured in RPMI with 10 % foetal bovine serum, 2 mM L-Glutamine and 100 µg/mL penicillin-streptomycin at 37 °C and 5 % CO_2_. MR49F was treated with 10 µM of enzalutamide to retain resistant phenotype.

Cells were grown in media supplemented with charcoal stripped foetal bovine serum, then treated in triplicate wells for 24 hours with either DMSO (0.5%; vehicle control simulating ADT-condition), DHT (1 nM; AR-agonist), enzalutamide (10 µM; AR-inhibitor) or DHT plus enzalutamide. Cells were then detached using 0.53 mM EDTA, fixed (1 % paraformaldehyde (PFA) (Electron Microscopy Services), 15 min), frozen overnight at –80 °C, thawed and spun onto 96-well glass-bottomed plate pre-coated with 3.5 µg/cm^2^ Cell-Tak.

### CTC enrichment

Blood samples were centrifuged at 300 g for 10 min, most of the plasma was collected and the cellular fraction resuspended in the remaining plasma plus PBS and added to a Sepmate tube (Stemcell Technologies) preloaded with 14 mL of Lymphoprep, then centrifuged at 1200 g for 10 min. After one PBS wash cells were resuspended in 8 mL PBS and CTC enrichment performed using the Genesis cell isolation system (BioRad) as per manufacturer’s instructions. Enriched cells were pelleted by centrifugation (at 300 g/10 mins) and supernatant was removed leaving 1 mL for resuspension after which 1 mL of 2% pathology grade PFA was added. After 10 mins the cells were pelleted and the pellet suspended in 1 mL of the patient’s plasma with 5% DMSO and immediately frozen at –80 °C using a Coolcell (Corning). For longer term storage samples were transferred to liquid nitrogen.

### Multiplexed imaging via Cyclic Immunofluorescence (CyclicIF)

#### Cyclic immunofluorescence labelling

96 well #1.5 glass bottom plates (Cellvis, CA) were pre-coated per well with 40 µL Cell-Tak at 3.5 µg/cm^2^ in NaHCO_3_ (pH 8) for two hours. Excess Cell-Tak was removed, wells were dried, washed twice with 200 µL H_2_O and dried again. Cryovials of enriched CTCs were thawed at 37 °C before immediate centrifugation at 400 g for 10 mins. Supernatant was removed and replaced with 200 µL of PBS; both without disturbing the pellet. Cells were centrifuged again at 400 g for 10 mins, supernatant was removed, and cells were resuspended in 200 µL of PBS and then added per well of the 96-well plate. Cryovials were washed with a further 100 µL of PBS that was then added to the same well. Plates were spun (300 g, 10 mins) to promote cell attachment. For each CycIF run, fixed, frozen DU145 cells from the same batch of a single passage were thawed and centrifuged onto a well for autofluorescence, bleaching and batch normalisation controls.

To limit cell loss, a minimal dead volume of 50 µL per well was maintained through all washing and labelling steps, with all subsequent reagents added at 2X concentrations in 50 µL for a final volume of 100 µL. All subsequent aspiration (50 µL) and washing (370 µL) steps were automated using the 50 TS microplate washer (BioTek), with an optimized washing procedure described in **Supplementary Fig. 1**. After centrifugation, wells were aspirated, and cells were fixed with fresh 4 % paraformaldehyde for 15 mins. Wells were aspirated again permeabilized with 0.1% Triton X-100 (Sigma-Aldrich) for 15 mins, after which wells were aspirated, and cells were fixed again to the plate with 4 % paraformaldehyde for 15 mins. Wells were washed three times, and cyclic immunofluorescence was commenced.

CycIF comprised of four repeated steps. First, cells were blocked with 4 % BSA in PBS for 15 mins. Secondly, wells were aspirated to minimum dead volume, and antibodies were added (names, fluorophores, sources and concentrations as per Table 2). As two CycIF cycles were conducted daily, antibodies were stratified based on their signal strength; high-affinity antibodies were incubated with rocking for 2 hours at room temperature, and weaker antibodies were incubated overnight at 4°C. After incubation, wells were aspirated and cells were labelled with DAPI at 1 µg/mL in 4 % BSA for 15 min. Wells were aspirated before washing three times with PBS. Prior to imaging, wells were filled to the top with PBS and covered with an ibidiSeal cover film (Ibidi) for consistent differential interference contrast (DIC) imaging. Fluorescence images were acquired (see Confocal Imaging below) after which wells were aspirated and bleaching solution (20 mM NaOH, 3 % H_2_O_2_ in PBS) was added and incubated under an LED table lamp (generating no heat) for one hour. Six PBS washes preceded the next CycIF cycle.

**Table 2:** Antibody reagents, fluorophores, catalogue numbers and usage concentrations.

| 488 | 555 | 647 |
| --- | --- | --- |
| p-MEK1, Cat #ab309797, RRID:AB_3675213, Concentration 1:500 | *PDL-1 (CD274), Cat #ab213358, RRID:AB_2728794, Concentration 1:500 | p-EGFR, Cat #ab205828, RRID:AB_2890267, Concentration 1:500 |
| CD276, Cat #ab218280, RRID:AB_3675218, Concentration 1:500 | Synaptophysin, Cat #ab206870, RRID:AB_205828, Concentration 1:500 | PTEN, Cat #ab310222, RRID:AB_3675215, Concentration 1:500 |
| *ALDH1A1, Cat #ab195254, RRID:AB_3675220, Concentration 1:500 | Lamin A+C, Cat #ab210433, RRID:AB_3073678, Concentration 1:500 | P-Jak1, Cat #ab311068, RRID:AB_3675219, Concentration 1:500 |
| NCAM-cd56, Cat #ab396808, RRID:AB_396808, Concentration 1:500 | CXCR4, Cat #ab211982, RRID:AB_3675221, Concentration 1:500 | YAP, Cat #ab225440, RRID:AB_3095991, Concentration 1:500 |
| HER-2, Cat #ICR55, RRID:AB_3100528, Concentration 1:500 | *TWIST, Cat #bs-2441R-A555, RRID:AB_3674791, Concentration 1:500 | p-SMAD2/3, Cat #562696, RRID:AB_2716578, Concentration 1:500 |
| Dlx1, Cat #bs-6467R-A488, RRID:AB_3674793, Concentration 1:500 | EpCaM, Cat #5488, RRID:AB_10693336, Concentration 1:500 | Vimentin, Cat #677807, RRID:AB_2616801, Concentration 1:500 |
| TGFBR2, Cat #bs-0117R-A488, RRID:AB_3674795, Concentration 1:500 | AR, Cat #8956, RRID:AB_11129223, Concentration 1:500 | CD146/MCAM, Cat #OJ79C, RRID:AB_770078, Concentration 1:500 |
| EGFR, Cat #5616, RRID:AB_10691853, Concentration 1:500 | pAkt (THR 308), Cat #77440, RRID:AB_3675230, Concentration 1:500 | PSA, Cat #bsm-1625M-A647, RRID:AB_3674790, Concentration 1:500 |
| STING, Cat #78827, RRID:AB_3675228, Concentration 1:500 | phospho-ERK, Cat #14095, RRID:AB_2728834, Concentration 1:500 | NF-κB, Cat #bs-1194R-A647, RRID:AB_3674789, Concentration 1:500 |
| p-STAT3(Tyr705) (clone DA37), Cat #4323, RRID:AB_561299, Concentration 1:500 | Ki67, Cat #20701, RRID:AB_3675231, Concentration 1:500 | p85/p55 PI3K, Cat #81913, RRID:AB_3675227, Concentration 1:500 |
| ERK, Cat #4780, RRID:AB_1903964, Concentration 1:500 | CD44, Cat #8724, RRID:AB_10829611, Concentration 1:500 | pAkt (Ser473), Cat #4075, RRID:AB_916029, Concentration 1:500 |
| c-KIT (CD117), Cat #3310, RRID:AB_823451, Concentration 1:500 | KRAS, Cat #64520, RRID:AB_3675236, Concentration 1:500 | mTOR, Cat #5048, RRID:AB_10828101, Concentration 1:500 |
| P53, Cat #2015, RRID:AB_2206297, Concentration 1:500 | *ACTIVE β-catenin, Cat #05-665-af555, RRID:AB_3675237, Concentration 1:500 | pSTAT5, Cat #9365, RRID:AB_1904151, Concentration 1:500 |
| p-GSK3B, Cat #14026, RRID:AB_2798365, Concentration 1:500 | *SLUG, Cat #bs-1382R-A555, RRID:AB_3674797, Concentration 1:500 | p-P53, Cat #8695, RRID:AB_10896847, Concentration 1:500 |
| cGAS, Cat #34256,<br>RRID:AB_2943225,<br>Concentration 1:500 | β-catenin total, Cat #71-2700,<br>RRID:AB_2533982,<br>Concentration 1:500 | Caspase-3 cleaved, Cat #9602,<br>RRID:AB_2687881,<br>Concentration 1:500 |
| pan-cytokeratin, Cat<br>#AE1/AE3,<br>RRID:AB_1834351,<br>Concentration 1:500 | ARV7, Cat #E308L,<br>RRID:AB_2798823,<br>Concentration 1:500<br>Custom conjugated | AKT, Cat #05-591-af647,<br>RRID:AB_3675238,<br>Concentration 1:500 |
| PSMA, Cat #GCP05,<br>RRID:AB_2539535,<br>Concentration 1:500 | C-MYC, Cat #AB201775,<br>RRID:AB_3675214,<br>Concentration 1:500 | CD45, Cat #H130,<br>RRID:AB_2802301,<br>Concentration 1:500 |
| Snail1, Cat #53-9859-82,<br>RRID:AB_2574483,<br>Concentration 1:500 | p-p38 MAPK, Cat #8632,<br>RRID:AB_2797648,<br>Concentration 1:500 | TRIB2D, Cat #bs-6262R-A647, RRID:AB_3674788,<br>Concentration 1:500 |

\*High background in patient samples.

#### Confocal Imaging

Confocal imaging was conducted on a Nikon AXR microscope using Nikon Jobs software for acquisition automation. Images were taken with a 20X PLAN APO 0.8 NA air-immersion objective, using a high-speed resonance scanner at a resolution of 2048 x 2048 with 8X line averaging. Channels were sequentially scanned for minimal crosstalk, with excitations of 405 nm, 488 nm, 555 nm and 647 nm generating emissions detected with bandpass filters of 429-474 nm, 499-530 nm, 571-625 nm and 662-737 nm, respectively. DIC images were also captured concurrent with 647 nm excitation.

Entire wells were scanned (45 image fields per well) to capture all cells per sample. For each well, the Nikon Perfect Focus System (PFS) was used for hardware (laser-based) autofocusing followed by fine-tuned image-based focusing using a Z-stack to determine strongest DAPI signal – defining the optimal Z-height for acquisition in each well. For acquisition, a Z-stack of 3 steps with 4 µm step-size was captured at each image field. This enabled subsequent Z-stack image max projection (see Image pre-processing below) to match focus for cells in non-flat fields. In each patient sample batch, DU145 cells from a single stored batch from a single passage were plated as consistent standards, and were then labelled and imaged with the same conditions as patient sample cells, guiding robust Z-score normalization of different patient batches (labelling as per **Supplementary Fig. 2**; normalization as per **Supplementary Fig. 3**).

Image files were saved as Nikon .nd2 files containing data per well, per CycIF cycle.

### Image Analysis

#### Image pre-processing and registration

Nikon .nd2 image files were exported as maximum projection .tiff files. Image registration was conducted using FIJI (ImageJ), beginning with the creation of a ‘registration’ channel for each CycIF cycle, using a smoothed DAPI image. Registration of DAPI images was then performed via the MultiStackReg FIJI plugin with a rigid body transformation. After registration, a minimum projection of DAPI images (from all CycIF cycles) was generated to exclude improperly registered cells, including cells that moved or were lost during CycIF cycles.

#### Single-cell segmentation and quantitative feature measurement

Segmentation and quantitative measurement of cells were conducted using Cell Profiler (4.2.6). Briefly, using minimum projections of registered DAPI images from all CycIF cycles, nuclear segmentation was performed. Cell body segmentation was constructed using the combined signals from α-tubulin, CD44, CD45, vimentin, HER-2, DAPI and mTOR. This mixture ensures that segmentation results are robust despite any variations in the expression level of a particular marker. Cell bodies contacting the image border were excluded to remove partial cells. For each remaining cell body, cytoplasm masks were determined by subtracting nuclear masks from cell body masks. Thereafter, for cell body, nucleus and cytoplasm segmentation, 78 area and shape features were measured per cell. Additionally, 156 texture features, 45 intensity features, 48 granularity features and 2 spatial ratio features (251 total features) were measured for each marker in each cell subdomain (cell body, nucleus, cytoplasm) (**Supplementary Table 1**).

#### Image and quantitative data integration, data parsing and data analysis

Quantitative cell data were parsed, processed and analysed using Knime 5.2^39^ (Zurich, Switzerland), with per cell quantitative features linked to images of corresponding cells. This integration allows for direct visual inspection and validation of per cell image and matched quantitative data. Poorly segmented and apoptotic cells were parsed via both quantitative selection and visual (image) inspection. All features from retained cells then underwent Z-score normalization ([per cell feature value – mean feature value] / feature standard deviation). Principal components analysis (PCA) was then performed for dimension reduction, retaining 97% of total dataset variance. Manifold embedding was then performed using principal components as inputs for uniform manifold approximation and projection (UMAP^40^). For color-coding of UMAP embeddings, cell features underwent either log_2_ or log_10_ transformations (depending on feature distributions) to improve color-scaling for visualization. To reconstruct progression in a progressive model of therapy resistance, pseudotime analysis was conducted as previously described^41^ using the ‘scFates’ Python package to model a cellular trajectory as a principal graph in low-dimensional UMAP-space. Specifically, a principal curve was fitted to the two-dimensional UMAP embedding, trending from sensitive (LNCaP) to resistant (MR49F) cell populations with hyperparameters set to: nodes=12; mu=10; lambda=0.01.

To perform cell-line or CTC classification, multiclass random-forest classifiers were used with well-holdouts to test model generalisability and robustness. For the cell line model, bootstrapping was performed to generate 2000 random samples, and Boruta feature selection was run iteratively on each sample to generate a list of features that were important in each run. Boruta identifies all features related to the target by comparing them to randomly shuffled shadow features, thereby making it suitable for marker discovery^42^. For CTC classification, the dataset was split 80/20 into train/test subsets. Prior to training, values underwent robust Z-score normalization as above, and under-sampling was applied to ensure balanced classes. Specifically, patient sample datasets were down-sampled 2000 times to generate 2000 class-balanced samples. All data visualisation was performed in Python 3.12 using seaborn and matplotlib packages, Boruta feature selection was run in R version 4.3.1.

## Results

### Multiplexed imaging data enables automated classification of PCa therapy-resistance states in a multi-stage resistance model

To validate the utility of deep multiplexed imaging for identifying disease progression and/or resistance drivers via longitudinal sampling of CTCs, we assessed a three-stage cell line model of PCa therapy resistance-progression as an established ground-truth exemplar. LNCaP cells (hormone-sensitive, enzalutamide-sensitive) were compared to two therapy-resistant, LNCaP-derived cell lines generated via xenograft modelling under *in-vivo* therapy exposure^37^. These are V16D cells (castrate-resistant, enzalutamide-sensitive) and MR49F cells (castrate-resistant, enzalutamide-resistant). With all cells grown in stimulus-depleted (charcoal-stripped) media, each cell line was then treated with either: DHT (1 nM, AR-agonist; simulating the presence of AR ligand in disease); DMSO (simulating ADT (AR ligand-depletion) in disease); DHT + ENZ (simulating AR-inhibition in the presence of AR ligands) or; Enzalutamide (10 µM; simulating AR-inhibition with ADT) for 24 hours, thus collectively emulating multiple phases of PCa treatment and resistance. Cells were then trypsinized and detached, fixed, frozen overnight, thawed and spun into a 96-well glass-bottomed plate pre-coated with Cell-Tak to enable non-specific cell attachment. These processing steps mimic all handling steps optimized for our liquid biopsy protocol post-CTC enrichment from blood samples (**Fig. 1A**). Once attached, cells underwent 14 sequential cycles of CycIF labelling, confocal imaging and fluorescence ablation to acquire a total of 42 protein and phospho-protein markers per cell (as well as DAPI and label-free differential interference contrast (DIC) channels, **Fig. 1A**). As detailed in methods, resulting image data were pre-processed (including illumination correction) and registered (to align all CycIF image cycles), permitting cell segmentation and single-cell quantitative feature measurement to delineate cell morphology and per marker expression intensities and subcellular localizations. Data were then parsed both quantitatively and via visual inspection to remove spurious objects, leaving a total of 4066 cells with detailed quantitative profiles spanning 10,707 features per cell.

**Figure 1:**
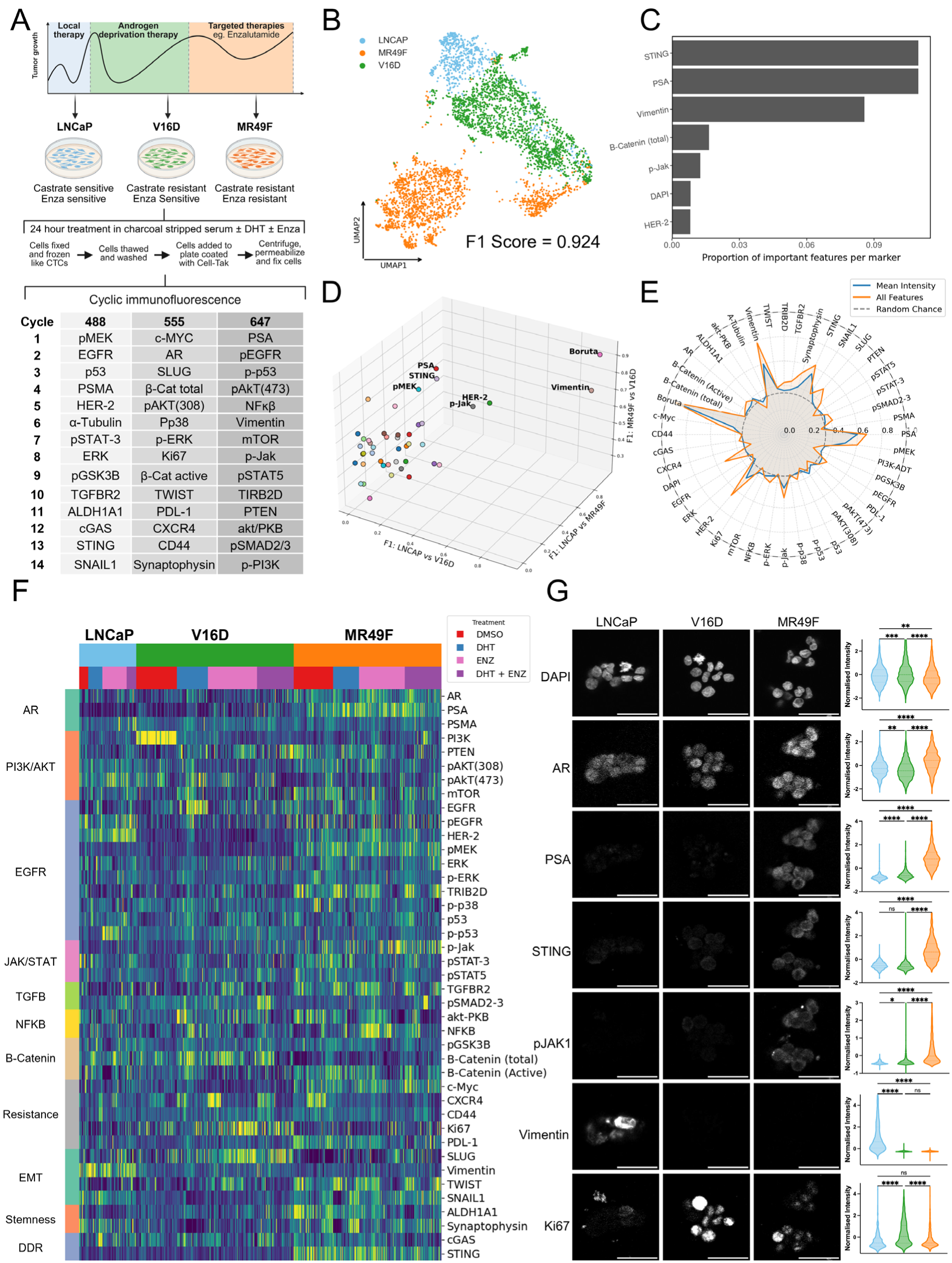
Deep multiplexed imaging delineates resistance states and associated marker expression in longitudinal model of PCa. (**A**) PCa cell lines LNCaP (castrate sensitive, n = 641) V16D (castrate resistant, n = 1766) and MR49F (castrate and enzalutamide resistant, n = 1649) previously derived via xenograft^37^ represent three different sensitivity states in the same cancer. These were treated with either DMSO (0.5%), DHT (1 nM), enzalutamide (ENZ, 10 µM) or DHT plus ENZ for 24 hours before being labelled via CyclicIF for the indicated molecular markers (n = 3 technical replicates per cell line, per treatment). (**B**) Per cell marker intensity feature-based UMAP shows clear clustering of color-coded cell lines. (**C**) Random forest machine learning classification identifies cells from each line with an average F1-score of 0.924. (**C**) Boruta feature selection bootstrapped over 2000 iterations ranked markers via importance for resistance classification, selecting 88 features (**Supplementary Table. 2**). Total feature importance of the top markers 7 was graphed. (**D**) Binary cell type classification F1-scores (LNCaP vs V16D, X axis; LNCaP vs MR49F, Y axis; V16D vs MR49F, Z axis) for each individual marker and optimised multivariate Boruta model (86 selected features). (**E**) F1-scores of random forest models classifying all cell types (concurrently) using per marker quantitative feature inputs including either: per cell mean intensity values only (blue line, 1 feature per marker), or; all per cell intensity and subcellular spatial features (orange line, 251 features per marker including intensity, granularity, texture and spatial ratio features); compared to random chance (grey dashed line). (**F**) Heatmap showing mean intensity of all markers in individual cells across cell types and treatments. (**G**) Representative images and mean intensity violin plots for LNCaP, V16D and MR49F cells labelled with DAPI, AR, PSA, STING, p-JAK1, vimentin and Ki67. Significance calculated via one-way ANOVA. Scale bars = 50 µm.

Unsupervised dimension reduction (via uniform manifold approximation and projection; UMAP^40^) of all features resulted in spontaneous clustering according to cell line identity, implying strong quantitative capacity to discern different therapy-resistance states with our deep multiplexed imaging data (**Fig. 1B**). MR49F cells further divided into two distinct clusters independent of experimental treatments. Automated random forest machine learning using all intensity, texture, and granularity features per marker successfully classified cells (from ‘hold-out’ cell subsets unseen during model training) according to the three resistance states associated with LNCaP, V16D and MR49F cell populations (average precision, recall, and F1-scores above 0.924 irrespective of experimental treatments, **Fig. 1B**). To identify the most important molecular marker features for discerning resistance states, robust Boruta feature selection^42^ was performed over 2000 iterations, with features selected in over 90% of iterations deemed as important (**Supplementary Fig. 2**). 86 features were identified (shown in **Supplementary Table 2**), with STING, prostate-specific antigen (PSA) and vimentin markers having the highest proportional representation among important features, along with features from β-catenin (Total), pJAK1, DAPI and HER-2 (**Fig. 1C**).

To differentiate whether these molecular markers are associated with the emergence of either castrate-or enzalutamide-resistance states, each marker was assessed on how they individually impact test performance of the random forest model. Vimentin had the highest performance for differentiating LNCaP from V16D and from MR49F (F1-scores of 0.91 and 0.93, respectively), whereas PSA and STING had the highest performance differentiating castrate-resistant V16D from castrate-resistant and enzalutamide-resistant MR49F (F1-scores 0.93 and 0.86, **Fig. 1D**). The collective 86 Boruta features identified in **Fig. 1C** had the highest performance across all comparisons (**Fig. 1D**).

Next, we examined the importance for resistance state classification of features defining the subcellular localisation of molecular markers, as opposed to marker levels-only per cell. For each marker, the performance of a random forest model trained using mean cell intensity-only per marker (1 feature) was compared to another random forest model trained using all intensity and subcellular localization features per marker (251 features including intensity, granularity, texture and distribution ratios). Most markers performed better than random chance (F1-score 0.3455) and in most cases, there was increased performance using all (spatial and intensity) features, particularly with high performing markers such as vimentin, PSA, STING, HER-2 and DAPI (**Fig. 1E**). While there was only a small increase in prediction performance using intensity and subcellular spatial features (relative to intensity-alone) in the Boruta-optimized multi-marker model, the model including subcellular spatial features achieved significantly improved classification confidence; a critical requirement for translational application of predictive models (**Supplementary Fig. 4**). The Boruta-optimized multi-marker, multi-feature model (F1-score 0.92) outperformed any single-marker, multi-feature (or mean intensity-only) model (vimentin F1-score 0.79). These results again highlight the increased efficacy of multiplexed immunofluorescence for identifying therapy-resistance PCa states (**Fig. 1E**). Moreover, the improved classification performance achieved by including intensity *and* subcellular spatial features (rather than mean intensity-only) highlights the importance of intracellular molecular marker localization in delineating cell resistance states. This in turn emphasizes the value for CTC profiling of multiplexed imaging as opposed to other multiplexed molecular sampling methods that lack subcellular resolving power, and aligns with evidence that subcellular localization functionally regulates most proteins^43^.

A heatmap of all cells (arranged by resistance state and treatment) with markers ordered by association with key signalling pathways or states reveals distinctions between, and heterogeneity within, each cell line, as well as co-expression patterns connecting various molecular markers (**Fig. 1F**). This detailed molecular mapping delineates numerous patterns linked to different stages of therapy response and resistance. For instance, LNCaP cells show increased p-p53, indicative of apoptosis, reflecting their sensitivity to enzalutamide treatment, while V16D cells under DMSO treatment (emulating ADT given charcoal-stripped media) show strong and ubiquitous p85(Tyr458)/p55(Tyr199) PI3K (p-PI3K) signalling, a major driver of ADT-resistance^44^. By contrast, V16D cells display increased EGFR expression^45^ under DHT stimulation of AR signalling (**Fig. 1F**).

To expand this analysis, we present example images of markers defined by machine learning as most discriminatory of resistance states, as well as key PCa markers, along with quantitative comparison (violin plots) of marker expression per cell line (**Fig. 1G**). In comparison to LNCaP, MR49F cells showed significantly increased AR localisation to the nucleus (P < 0.0001, one-way ANOVA), indicating increased activity, which is supported by increased expression of AR’s downstream target, PSA (P < 0.0001); a well-defined dependence^46^ (**Fig. 1G**). In comparison to V16D, MR49F also showed increased expression of STING (P < 0.0001), as well as increased phosphorylation of both JAK1 (pJAK1) and MEK (pMEK) (both P < 0.0001), again corroborating previous reports of mechanisms linked to enzalutamide-resistance^47,48^. LNCaP cells on the other hand displayed significantly higher vimentin expression, which was lost in both V16D and MR49F cells (both P < 0.0001). Compared to both LNCaP and MR49F, V16D cells expressed significantly increased levels of the proliferation marker, Ki67 (both P < 0.0001, **Fig. 1G**).

### Multiplexed image data enables pseudotemporal inference of dynamic transitions between resistance states

Having determined that multiplexed imaging data can *discretely* classify PCa therapy-resistant cell states, we next interrogated *continuous* changes in molecular expression levels (mean intensities per marker per cell) associated with transitions through resistance states. Unsupervised pseudotime trajectory inference^41^ delineated a path spanning LNCaP via V16D to MR49F cells through UMAP-defined state space (**Fig. 2A**). With each cell assigned a pseudotime value (**Fig. 2B**) and cell type-specific population densities plotted *versus* pseudotime (**Fig. 2C, left**), average mean cell intensities (for each of 22 pseudotime value bins) per multiplexed marker were plotted against pseudotime and hierarchically clustered to reveal an array of complex yet progressive dynamics (**Fig. 2C, right**). Initially, vimentin, α-Tubulin, HER-2 and SNAIL1 were highly expressed in LNCaP cells before rapid reductions. Transition to V16D corresponded with increased β-catenin and p-PI3K levels, and a steady increase in proliferative and epithelial to mesenchymal transition (EMT) markers Ki67 and SLUG (**Fig. 2C**). With transition to the enzalutamide resistance of MR49F populations around pseudotime value 8, there was a reduction in most markers, which coincided with low DAPI indicating that this population may be in cell cycle phase G0/G1. Subsequently, a sharp increase was observed in PSA, STING, pMEK and pJAK1, along with increased expression of various signalling, EMT, stemness and DNA damage-response (DDR) markers (**Fig. 2C**). AR expression was also increased in enzalutamide-resistant MR49F cells with high pseudotime values.

**Figure 2:**
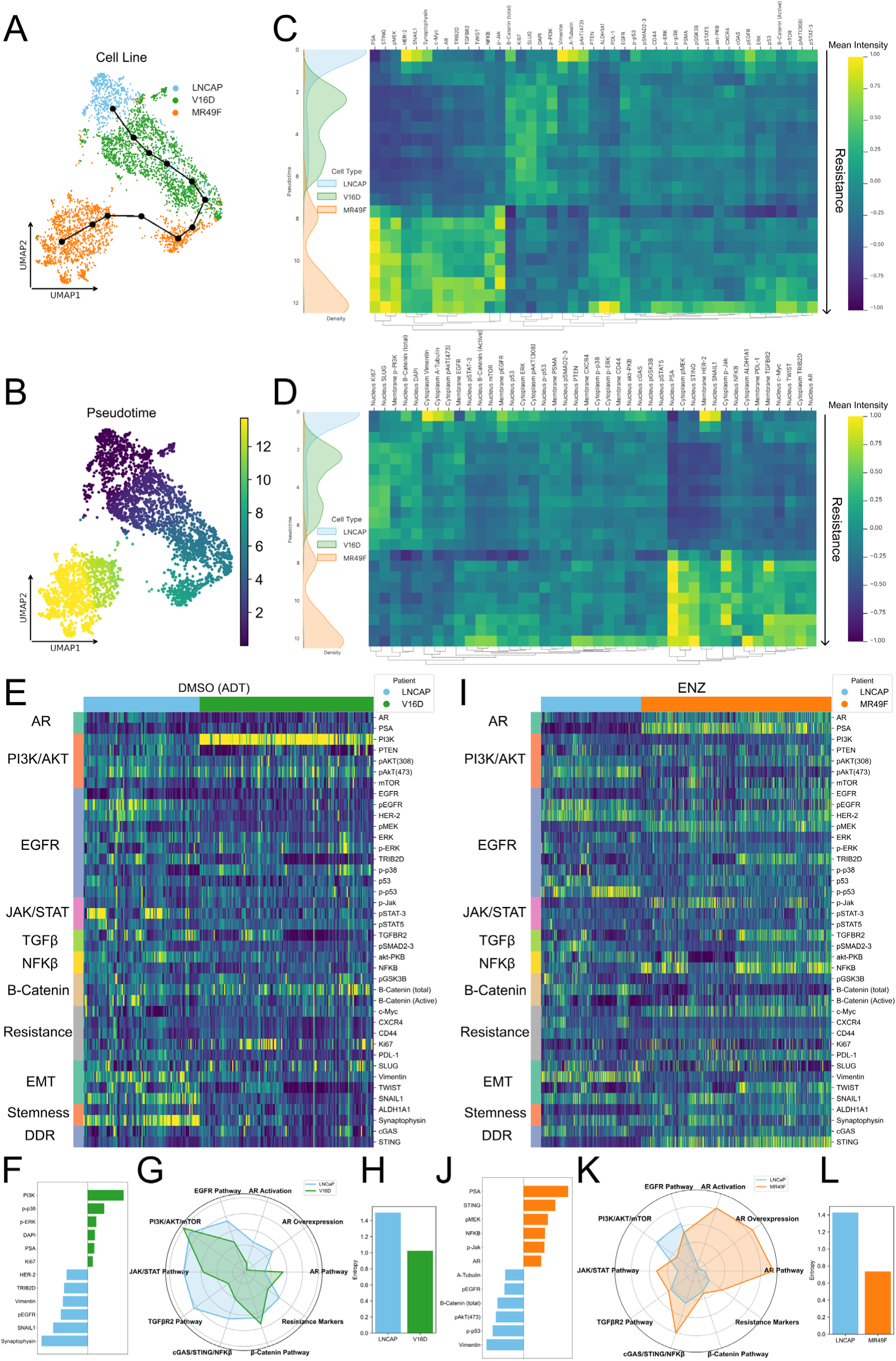
Deep multiplexed imaging identifies protein subcellular localisation and pathway activation correlating with the progressive emergence of resistance. (**A**) Unsupervised pseudotime trajectory plotted through UMAP manifold (as per **Fig. 1B**) from LNCaP (sensitive, blue) through V16D (castrate-resistant, green) to MR49F (enzalutamide-resistant, orange) using scFates^41^. Pseudotime values assigned per cell (**B**) given trajectory in A. Heatmaps of (**C**) mean cellular (**D**) and membrane, cytoplasmic or nuclear marker intensities plotted against pseudotime with density distributions of corresponding cell lines plotted (left). (**E**) Heatmap (arranged by cell type, X axis; markers and marker categories, Y axis) and (**F**) waterfall plot of the top six positive and negative differences in marker expression between LNCaP and V16D cells under DMSO treatment. (**G**) Radar plot of marker intensities from key signalling pathway categories indicates activation of pathways known to confer resistance to ADT in LNCaP and V16D. (**H**) Shannon’s entropy index for LNCaP versus V16D cell populations. (**I**) Heatmap and (**J**) waterfall plot of the top six positive and negative differences in marker expression between LNCaP and MR49F under enzalutamide treatment. (**K**) Radar plot of marker intensities from key signalling pathway categories indicating pathway activation profiles. (**L**) Relative entropy of LNCaP versus MR49F cell populations. Signalling pathway categories used in radar plots defined in **Supplementary Table 3**.

Next, we leveraged the subcellular resolution of our image-based profiling by assessing mean marker intensities at specific subcellular localizations with known relevance to the activity of each marker (e.g. receptor expression at the plasma membrane, phospho-signalling markers in the cytoplasm, transcription factors in the nucleus). Similarly to **Fig. 2C**, these localized intensities were plotted via a heatmap relative to the pseudotime trajectory inferring changes in resistance (**Fig. 2D**), accentuating subcellular marker dynamics with functional implications. We thus detected high levels of nuclear SNAIL and synaptophysin in LNCaP cells, followed by increased nuclear SLUG, Ki67 and β-catenin spanning V16D cells, transitioning to heightened nuclear PSA, p-JAK1, STING and p-MEK in enzalutamide-resistant MR49F cells (**Fig. 2D**). Transcription factors including AR, c-Myc, TWIST and TRIB2D also showed increased nuclear accumulation in MR49F cells, implying their enhanced activation in the enzalutamide-resistant phase (**Fig. 2D**). Receptors including HER2 and phosphorylated EGFR displayed rapid declines in membrane labelling in LNCaP cells, as did cytoplasmic vimentin and α-tubulin, while PD-L1 membrane levels reached their peak at the highest pseudotime levels in MR49F cells, echoing reports of increased PD-L1 expression in enzalutamide resistance^49^ (**Fig. 2D**). These results illustrate the capacity of multiplexed imaging to reveal complex subcellular molecular dynamics with strong functional implications.

### Comparison of resistance states under therapeutic conditions identifies known castration– and enzalutamide-resistance drivers

To specifically investigate the emergence of castration resistance, LNCaP and V16D cells were compared under DMSO treatment in charcoal-stripped (i.e. hormone-depleted) media, thereby mimicking the ADT condition given the absence of AR-ligand DHT. A heatmap of mean intensity values per marker, per cell, shows that V16D cells displayed significant and near-uniform upregulation of p-PI3K relative to the castrate-sensitive LNCaP cell line, as well as reductions in the expression of synaptophysin, SNAIL1, Vimentin and pEGFR, among other markers (**Fig. 2E**). This highlights pathways with known associations to castrate resistance and increased proliferation, as identified by heightened Ki67^50^. A waterfall plot of average mean intensity marker levels comparing LNCaP *versus* V16D cells shows that p-PI3K is the most overexpressed feature in V16D, with downstream signalling proteins p-p38 and p-ERK also increased (**Fig. 2F**), and most EGFR-related markers decreased along with reduced stemness and mesenchymal markers (**Fig. 2F**). A radar plot combining marker intensities into signalling pathways categories (categories detailed in **Supplementary Table 3**) known to confer resistance to AR inhibition^51–56^ reveals reductions in signalling across most pathways except for the β-catenin and AR pathways (**Fig. 2G**). Notably, AR signalling pathway-status did not reveal either AR over-expression (cell mean intensity) or activation (nuclei intensity) at this intermediate resistance stage. Instead, PI3K signalling was highlighted as a central driver of castrate resistance in V16D cells, as has been reported in CRPC^50,57^. The molecular heterogeneity present within each cell population was quantified via Shannon’s entropy index, indicating reduced entropy (lower heterogeneity) across the V16D cell population (**Fig. 2H**). This relative homogeneity may reflect the effects of selection associated with the emergence of acquired castrate resistance.

To next interrogate enzalutamide resistance, LNCaP and MR49F cells were compared under enzalutamide treatment. A heatmap of mean intensity values per marker (**Fig. 2I**) and waterfall plot of maximal marker-level differences (**Fig. 2J**) highlight a substantial population of LNCaP (enzalutamide-sensitive) cells with strong p-p53 signalling, suggestive of stress and potential apoptosis-induction under enzalutamide treatment. This stress response was supported by high pAKT(S473) and vimentin levels, which are linked to enzalutamide-induced oxidative stress^58–60^ (**Fig. 2I-J**). PSA increases were most pronounced in MR49F cells, implying heightened AR signalling. Indeed, the most striking signature of enzalutamide resistance in MR49F cells was increased AR pathway signalling, denoted by both increased AR expression and nuclear localization (**Fig. 2K**). This corresponds to reports that MR49F cells have a full AR gene amplification compared to LNCaP cells, as well as frequent AR F877L activating mutations^61,62^. STING and NF-κB expression was also increased in MR49F cells, with STING known to activate NF-κB signalling, which in turn promotes enzalutamide resistance^63^. MR49F cells also exhibited increased p-MEK and p-JAK1 signals, as well as increased expression of c-MYC, PD-L1 and TRIB2D^64–66^, established PCa drivers and mediators of enzalutamide resistance. Ultimately, we detect a constellation of upregulated markers defining signalling pathways (AR, cGAS-STING-NF-κB and JAK/STAT signalling) and additional PCa drivers that are all strongly linked to enzalutamide resistance^47^. Like the V16D population, the MR49F cell population displays reduced entropy compared to LNCaP, with this relative homogeneity again suggestive of a selected phenotype (**Fig. 2L**).

### Deep multiplexed imaging of PCa liquid biopsy-derived CTCs enhances CTC identification

Standard ∼9 ml peripheral blood draws were collected constituting liquid biopsy samples from 7 PCa patients undergoing treatment with various combinations of ADT, enzalutamide (ENZ) and docetaxel (**Table 2**). By combining expert visual classification of CTCs with quantitative data parsing, high-confidence CTCs were detected in blood from 5 patients, 3 of whom were clinically classified as therapy ‘responders’ (patient 2, 5, 6) and 2 as ‘non-responders’ (patient 3, 4), based on their PSA trajectories (**Fig. 3A**) and clinical presentation. CTCs were enriched via size exclusion, before undergoing 16 CyclicIF cycles to characterise up to 50 markers per cell (**Fig. 3B**). Cell segmentation and labelling for an example CTC and immune cell is demonstrated (**Fig. 3C**).

**Figure 3:**
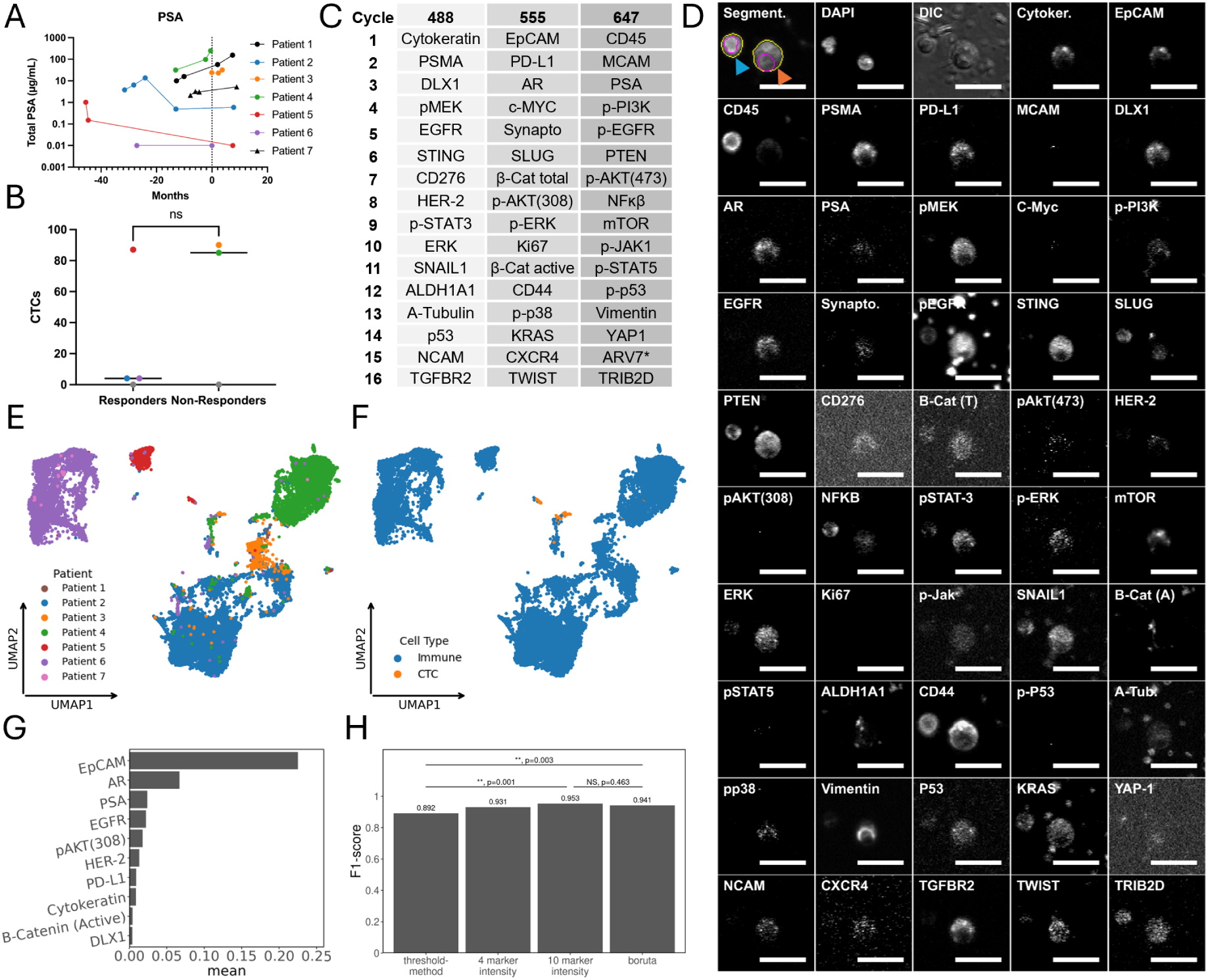
Deep multiplexed imaging improves CTC identification. CTCs from patients (n = 7) in Table 2 were isolated via size-exclusion and labelled with 16 cycles via CycIF. 5 patients had identified CTCs, of which responders were patients 2, 5 and 6 (blue, red and purple, respectively) and non-responders were patients 3 and 4 (orange and green, respectively). (**A**) PSA trajectories of patients with collection of CTCs at timepoint 0. (**B**) Enriched CTC populations were labelled with 16 cycles via CycIF (*ARv7 not labelled in all patients thus removed). (**C**) Example CTC (orange) and immune cell (blue) from patient 2 showing segmentation and labelling for all markers used in data analysis. (**D**) CTC numbers detected from responders and non-responders (**E**) Per cell marker intensity feature-based UMAP of all collected cells (n = 22, 192 immune cells, n = 270 CTCs) showing patient heterogeneity and (**F**) clustering of CTCs (orange) and immune cell (blue) populations. UMAP of all collected CTCs generated from intensity features. (**G**) Markers with the highest proportion of important features after 2000 iterations of boruta feature selection. (**H**) F1-scores comparing CTC identification performance using: canonical stepwise-thresholding CTC definition (DAPI positive, cytokeratin positive, CD45 negative); random forest classification of CTCs using intensity features from 4 canonical markers (DAPI, cytokeratin, EpCAM and CD45); random forest classification using intensity features from the top 10 markers identified in (**3G)** or; from the 22 Boruta features described in **Supplementary Figure 5**.

**Table 2:**
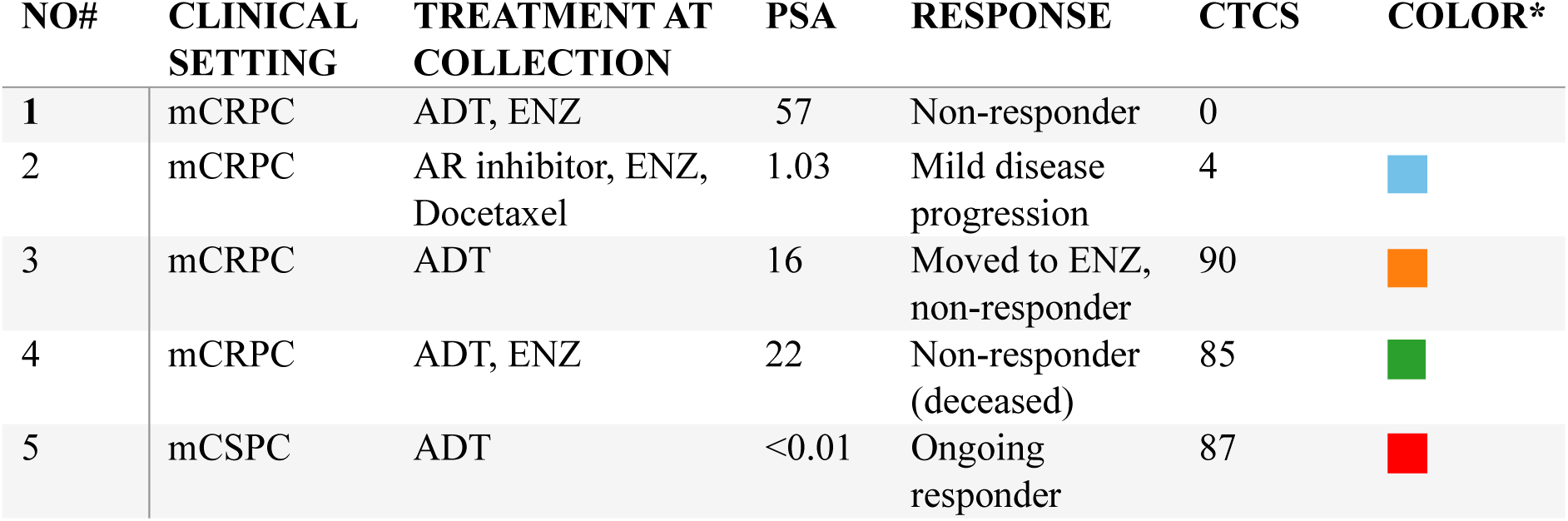
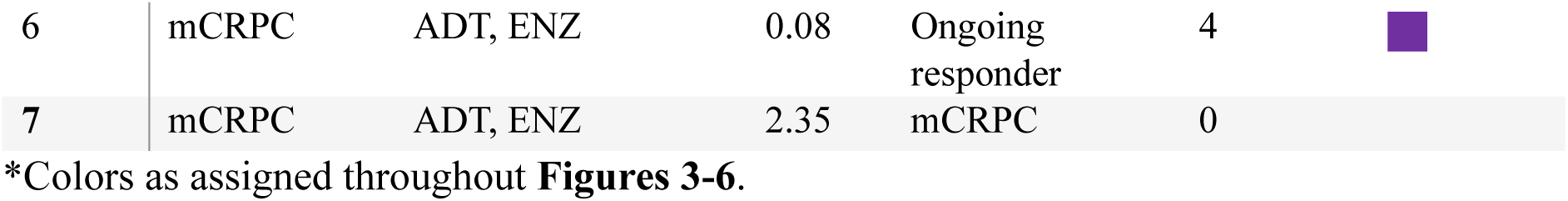
Patient number, clinical setting, treatment, blood PSA levels and clinically defined response of patients recruited for CTC analysis.

| NO# | CLINICAL SETTING | TREATMENT AT COLLECTION | PSA | RESPONSE | CTCS | COLOR* |
| --- | --- | --- | --- | --- | --- | --- |
| 1 | mCRPC | ADT, ENZ | 57 | Non-responder | 0 |  |
| 2 | mCRPC | AR inhibitor, ENZ, Docetaxel | 1.03 | Mild disease progression | 4 |  |
| 3 | mCRPC | ADT | 16 | Moved to ENZ, non-responder | 90 |  |
| 4 | mCRPC | ADT, ENZ | 22 | Non-responder (deceased) | 85 |  |
| 5 | mCSPC | ADT | <0.01 | Ongoing responder | 87 |  |
| 6 | mCRPC | ADT, ENZ | 0.08 | Ongoing responder | 4 |  |
| 7 | mCRPC | ADT, ENZ | 2.35 | mCRPC | 0 |  |

\*Colors as assigned throughout **Figures 3-6**.

This yielded in total 22,192 non-CTC residual blood cells and 270 CTCs across all patients. No significant difference in CTC counts was observed between responders and non-responders (P = 0.6571, Mann-Whitney test, **Fig. 3D**). After robust z-score-based batch-normalization of all quantitative features relative to concurrent labelling and imaging of DU145 control PCa cells (stored fixed and frozen in aliquots from a single passage, **Supplementary Fig. 2 & 3**), UMAP clustering was performed using all (immune and CTC) cells based on all intensity features per marker, per cell. Comparing all cells between patients, we observed only modest overlap with large-scale differences largely corresponding with patient identity (**Fig. 3E**). Comparing between immune and CTCs, we note relatively stronger overlap between CTCs, which cluster centrally within the UMAP and are distinct from the majority of immune cells and their largely patient-specific clusters (**Fig. 3F**).

CTCs are canonically defined as being DAPI-positive, EpCAM and/or cytokeratin-positive, and CD45-negative^67^. While these are useful markers for CTC detection from most solid carcinomas, CTCs often have lower expression of cytokeratin and EpCAM than cell lines^68^ and this expression can be lowered or lost in cells that have undergone partial or complete EMT^69^. To explore alternative approaches for CTC identification given our highly multiplexed datasets, we assessed whether machine learning-based CTC detection (i.e. CTC classification) can be improved by using different marker combinations. Using manually defined ‘ground-truth’ CTC identities derived via expert visual inspection using all markers, Boruta feature selection was conducted using all intensity features to determine which markers best predicted CTC identity. The 10 markers with the highest proportion of important features were EpCAM (0.225), AR (0.067), PSA (0.024), EGFR (0.022), pAKT(308) (0.018), HER-2 (0.013), PD-L1 (0.009), Cytokeratin (0.009), active β-catenin (0.004) and DLX1 (0.004) (**Fig. 3G**). A second round of Boruta feature selection was conducted using all intensity and subcellular spatial features (texture, granularity etc). 22 features were selected in over 90% of (2000) training iterations, and all of these were from either EpCAM (16) or AR (6). This highlights these two markers as invaluable for CTC identification in the PCa patient samples here assessed (**Supplementary Fig. 5**).

To compare our machine learning-based approach to a more canonical method of CTC identification, we assessed stepwise intensity thresholding-based definition^51^ of DAPI-positive, cytokeratin/EpCAM-positive and CD45-negative CTCs using empirically determined thresholds, resulting in an F1-score (relative to manual CTC definition) of just 0.8920 (**Fig. 3H**). Random forest machine learning-based CTC classification showed better performance, when using intensity features of either: the 4 canonical CTC markers (DAPI, CD45, cytokeratin and EpCAM, 180 features, F1-score 0.931); or the top 10 markers from **Fig. 3G** (450 features, F1-score 0.953); or the 22 Boruta-selected features from **Supplementary Fig. 5** (F1-score 0.941, **Fig. 3H**). Both the 10-marker and Boruta models performed significantly better than the sequential thresholding method typically used for CTC identification (p = 0.001 and 0.003, respectively, bootstrap Wald’s test), with no significant difference between these two machine learning models (p=0.463, **Fig. 3H**).

### Multi-molecular expression and signalling signatures hint at the discernability of therapy-responsive and non-responsive PCa patients

A heatmap of all CTCs hierarchically clustered by mean marker intensity (**Fig. 4A**) highlights the propensity of CTCs from the same patient to self-cluster, revealing overarching marker homologies. For example, CTCs from non-responder patients 3 and 4 have the highest PSMA. Notably, at the time of blood sampling, patient 4 was non-responsive to ongoing ADT and enzalutamide treatment, while patient 3 was not yet receiving enzalutamide but subsequently showed no clinical response when provided enzalutamide therapy. By contrast, CTCs from therapy-responsive patient 5 show high EGFR/PI3K/AKT expression and signalling (**Fig. 4A**). Importantly, this is similar to signalling profiles detected in the cultured LNCaP cell model of therapy-sensitive PCa under enzalutamide treatment (**Fig. 2J**).

**Figure 4:**
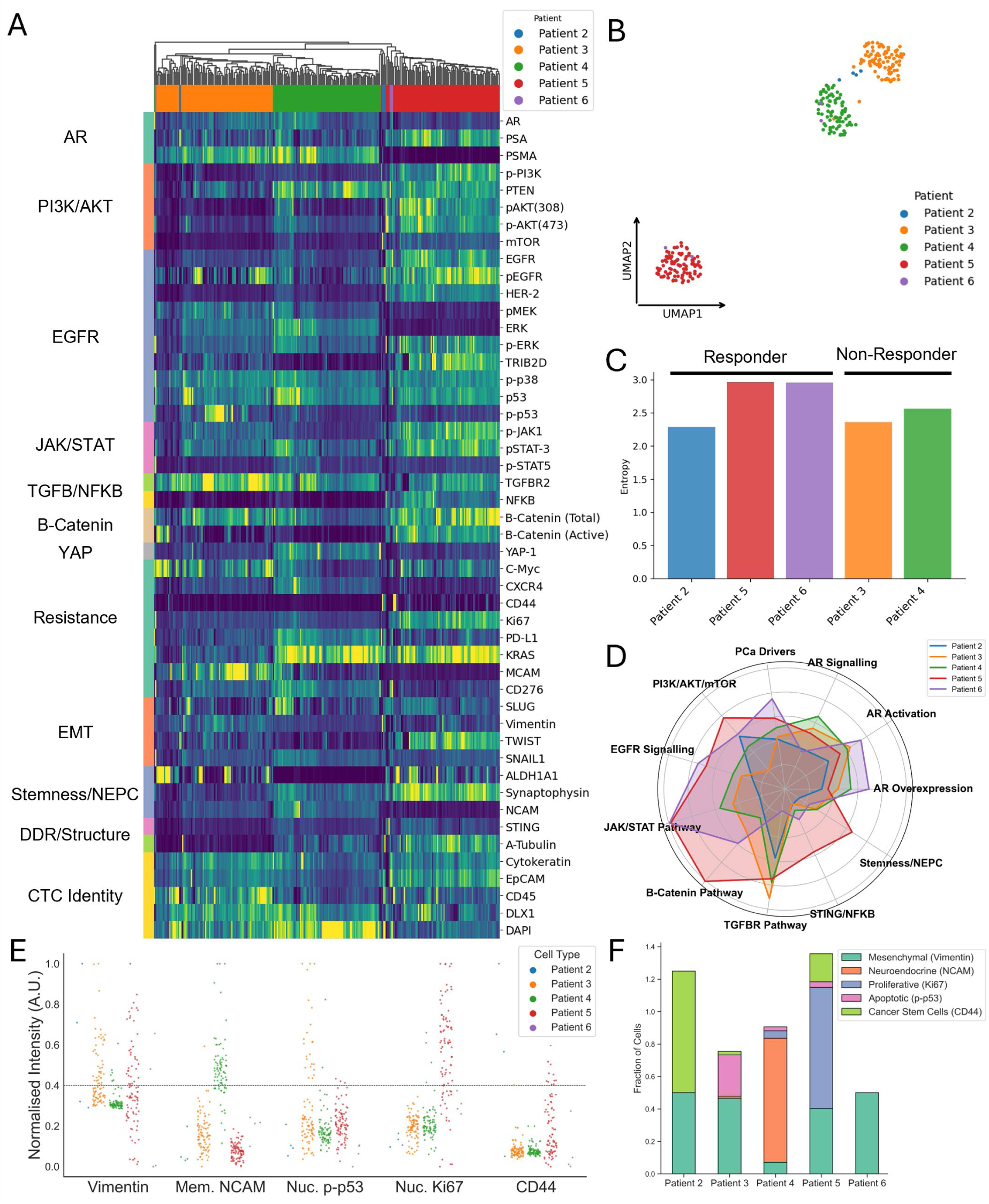
Deep multiplexed imaging allows single-cell characterization of PCa patient CTCs and identification therapy-response associated signalling pathway signatures. (**A**) Hierarchically clustered heatmap of mean marker intensities per CTC derived from blood-based liquid biopsy samples from PCa patients (color-coded on X axis, arranged by signalling pathway categories (Y axis). (**B**) Per cell marker intensity feature-based UMAP of CTCs (only) color-coded by patient number. **(C**) Shannon’s entropy index of variability in per patient CTC populations, quantifying relative heterogeneity. (**D**) Radar plot of mean pathway activation per patient. Signalling pathway categories used in radar plot defined in **Supplementary Table 3**. (**E**) A threshold of normalised mean intensity (rescaled from 0-99 percentiles) at 0.4 of key cell state markers of vimentin, membrane NCAM, nuclear p-p53, nuclear Ki67 and CD44 allows (**F**) phenotypic characterization of CTCs as mesenchymal, neuroendocrine, proliferative, apoptotic or cancer stem-cells, respectively. Phenotypes are not mutually exclusive, resulting in some fractions of CTCs being higher than 1.0.

Construction of a UMAP-embedding containing CTCs only revealed some overlap between patient cells, though here again CTCs primarily clustered by patient identity (**Fig. 4B**). Interestingly, this UMAP representation predominantly separates castrate-resistant prostate cancers from castrate-sensitive (patient 5). Patient 6 (with 4 CTCs detected) has two CTCs associated with the putative resistant phenotype (clustering with CTCs from patient 4), and two CTCs with the responsive phenotype (clustering with CTCs from patient 5) (**Fig. 4B**).

To further characterise CTC heterogeneity, Shannon’s entropy index was calculated for each patient’s CTCs, since this has been identified as a critical indicator of potential responsiveness to treatment^70^, including in our earlier comparison of resistant and non-resistant PCa cancer cell lines (**Fig. 2**). We found that CTCs from therapy-responsive patients 5 and 6 displayed the highest entropy values (**Fig. 4C**), aligning the therapy-sensitivity of these cancer cell populations with their heterogeneity, as also observed for the therapy-sensitive LNCaP cell line (**Fig. 2H & L**). By comparison, the therapy-non-responsive patients 3 and 4 showed lower CTC population-entropy, aligning their apparent therapy-resistance and reduced entropy with the same patterns seen in therapy resistance V16D and MR49F cells (**Fig. 2H & L**). As in the cell line model, this reduced entropy may reflect relative homogeneity imposed by selection of resistant cancer cell subpopulations. A radar plot of per patient CTC population mean signalling pathway signatures across various key categories (detailed in **Supplementary Table 3**) with known links to PCa therapy-responsiveness serves to define patterns of signalling activity in each patient (**Fig. 4D**). Different activity signatures are discernible between non-responsive (3 and 4) and responsive (5 and 6) patients. In fact, CTCs from responder patients 5 and 6 showed quite varied activity profiles, though these did include heightened PI3K/AKT, EGFR, JAK/STAT and β-catenin pathways (**Fig. 4D**; potentially indicative of stress-responses to therapy). The mixed signalling profiles in these CTCs mirror their high entropy (low uniformity). By contrast, CTCs from therapy non-responsive patients showed consistent and pronounced activation of the TGFβR pathway, a known driver of EMT and therapy-resistance in PCa CTCs^71,72^. Interestingly, we note that CTCs from therapy-responsive patient 2 clustered with CTCs from non-responsive patients 3 and 4 (**Fig. 4B**), likewise showing low entropy (**Fig. 4C**) and a similar increase in TGFβR signalling activity (**Fig. 4D**). While patient 2 is defined as therapy-responsive due to reduced blood PSA post-therapy, they are the only patient also receiving docetaxel. As their residual PSA levels remain ∼100 times higher than those of therapy-responsive patients 5 and 6, similarity in the molecular profiles of patient 2 and non-responder patients 3 and 4 may be indicative of either low response to ADT and ENZ, or emerging resistance in patient 2.

In addition to molecular scale biomarker expression and pathway activation, deep multiplexed imaging also reveals cellular-scale phenotypes of CTCs. An arbitrary but discriminatory threshold of 0.4 was manually selected and applied to the normalized mean intensities of state markers vimentin, membrane NCAM and CD44 (**Fig. 4E**), with CTCs above this threshold classified as mesenchymal, neuroendocrine or cancer stem cells, respectively (**Fig. 4F**). The same threshold was applied for mean nuclear intensities of Ki67 and p-p53, functioning as indicative biomarkers for proliferative and apoptotic phenotypes, respectively (**Fig. 4E-F**). CTCs from patient 2 were thus predominantly (75%) defined as cancer stem cells, whereas patient 3 had a sub-population (25.6%) of apoptotic CTCs, which may indicate a sub-population of CTCs susceptible (i.e. responsive) to treatment (**Fig. 4F**). 76.5% of patient 4 CTCs indicated neuroendocrine differentiation – an aggressive, enzalutamide-resistant phenotype well-characterized to have poorer outcomes^73^. Patient 5 showed a high proportion (74.7%) of proliferative CTCs, potentially an effect of its active EGFR/PI3K/JAK signalling (**Fig. 4F**).

### Subcellular molecular localization within CTCs highlights active and actionable markers

We next leveraged the subcellular spatial resolution provided by multiplexed imaging to assess whether intracellular marker localisations within CTCs provide additional information with potential utility. For simplicity, subcellular spatial distributions are here summarised in a hierarchically clustered heatmap of CTCs from all patients. The heatmap incorporates data on marker intensities at spatial domains related to known functions (per marker), including the: cell edge / plasma membrane (receptors, membrane proteins); cytoplasm (phospho-signalling proteins, kinases, inactive transcription factors, structural proteins) and; nucleus (active transcription factors) (**Fig. 5A**). The mean subcellular marker intensity for all CTCs from each patient is compared to the cohort mean, with the 10 largest deviations circled in black (**Fig. 5B**), revealing patient-specific, subcellularly localized marker enrichment. For instance, CTCs from patient 2 displayed high membrane CD44, cytoplasmic mTOR and vimentin as well as increased nuclear SNAIL1, SLUG and TWIST, indicating a mesenchymal phenotype, as well as low nuclear p53 levels. CTCs from patient 3 showed reduced plasma membrane KRAS, low cytoplasmic PTEN and EGFR-related markers. Patient 4 CTCs had very high membrane PSMA and KRAS expression. Conversely, patient 5 CTCs had undetectable PSMA expression, combined with active EGFR signalling, high cytoplasmic NF-κB, pJAK1 and high nuclear Ki67 levels. Similarly, patient 6 CTCs revealed very high EGFR expression and active EGFR signalling parameters detectable at membranes, cytoplasm and nucleus. CTCs from patients 3, 4 and 6 had the highest nuclear AR, highlighting its activation, while, interestingly, CTCs from patients 3 and 4 had detectable membrane CD45 expression, which has been previously reported to shield CTCs from immune surveillance^22^.

**Figure 5:**
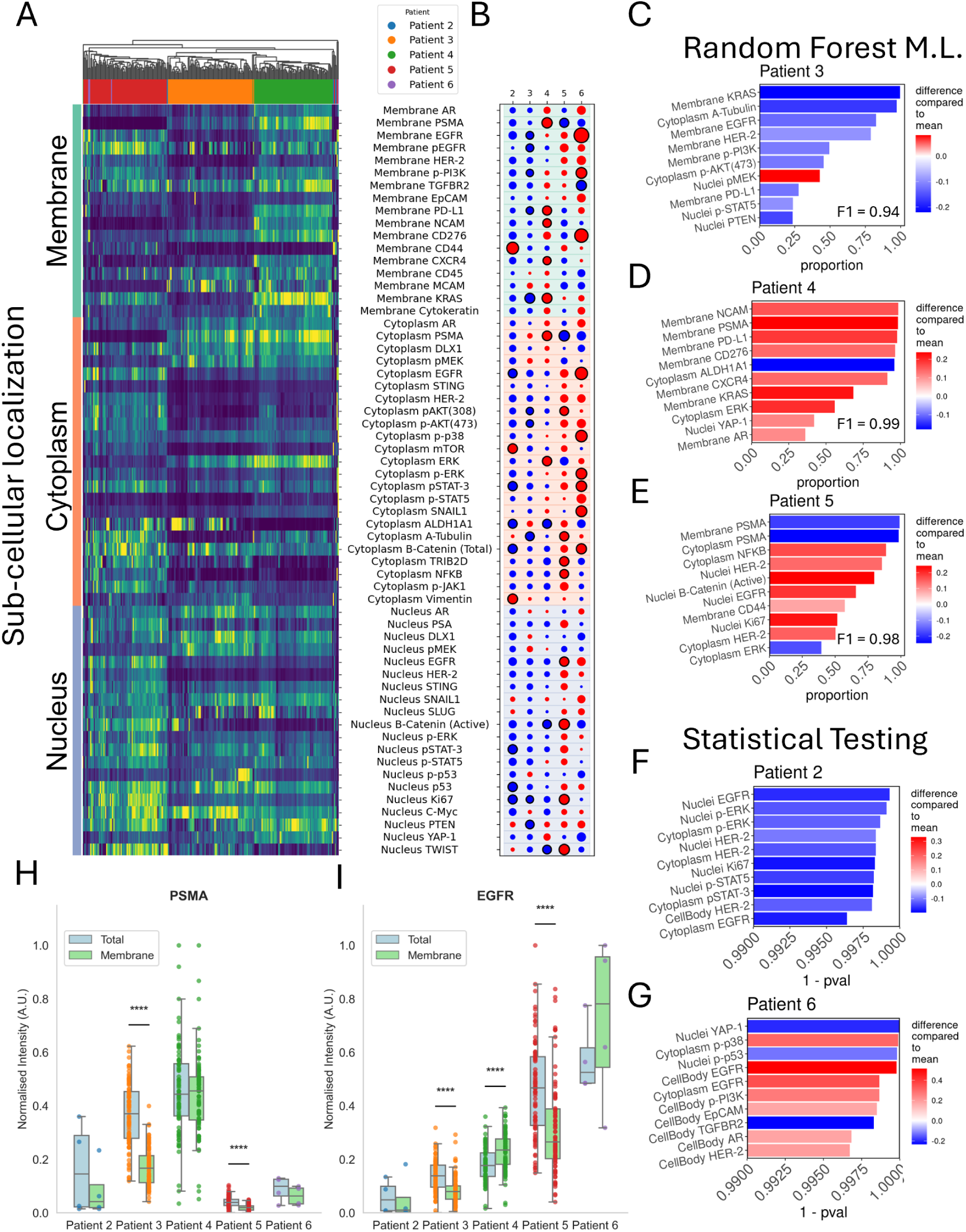
Deep multiplexed imaging enables detection of subcellular protein localisations delineating activated signalling pathways and distinct cell phenotypes. (**A**) Hierarchically clustered heatmap of per CTC marker intensities localized at the: cell edge (plasma membrane, green); cytoplasm (orange) or; nucleus (blue). (**B**) A bubble plot of mean subcellular intensities per marker showing higher expressions (red) and lower expression (blue) relative to the whole cohort mean. The largest 10 deviations from cohort mean per patient are circled in black. Random forest classification and feature selection were performed using per CTC subcellular marker localizations to delineate the top 10 most informative features (blue and red correspond to decreased or increased marker localization levels, respectively) distinguishing patient 3 (**C**) or patient 4 (**D**) or patient 5 (**E**) from the whole cohort. For patients 2 (**F**) and 6 (**G**) with lower CTC counts, the top 10 markers were identified according to p-values defined by the non-parametric Mann-Whitney statistical test. (**H**) Pairwise comparisons (Mann-Whitney test) of normalised PSMA and EGFR (**I**) total (blue) and membrane (green) intensities per patient.

To further identify markers and marker localizations that are distinctive between the CTC populations of each patient, random forest machine learning was run to classify CTCs from patients 3, 4 and 5. These patients were selected as their CTC numbers are sufficient for robust machine learning. This feature selection revealed the top 10 marker localizations delineating CTCs from each patient in comparison to the rest of the cohort. Therapy non-responsive patient 3 had CTCs with low KRAS, low α-tubulin and low p-PI3K signalling pathway activation, but high pMEK activation (**Fig 5C).** Therapy non-responsive patient 4 had CTCs with high membrane PSMA, as well as a striking neuroendocrine phenotype with increased expression of NCAM, CD276 and CXCR4 (**Fig. 5D**). Therapy responsive patient 5 had CTCs with very low expression of membrane PSMA, as well as high cytoplasmic NF-κB and high nuclear HER-2, EGFR, active β-catenin and Ki67 (**Fig 5E)**. Since four CTCs are not enough to conduct robust machine learning, feature selection for patients 2 and 6 was conducted using direct statistical methods. A binary comparison was performed for each biomarker between the CTCs of each patient and the remainder of the cohort, using the non-parametric Mann-Whitney test. Patient 2 had significantly lower expression of a range of markers in the EGFR signalling pathway (**Fig. 5F**), while patient 6 had significantly lower nuclear YAP-1 and p-p53, with very high membrane EGFR and cytoplasmic p-p38 signal (**Fig 5G)**.

The potential impact of sub-cellular marker localizations on precision oncology-relevant interpretations of CTC biology is showcased by pairwise comparisons of both PSMA and EGFR total *versus* membrane expression (**Fig. 5H-I**). While CTCs from patient 3 had a high total PSMA expression, these CTCs exhibited significantly lower (P < 0.0001, Mann-Whitney test) membrane intensity. This might imply reduced access and therefore efficacy for PSMA ligand-based diagnostics and therapeutics, which require surface PSMA expression (**Fig. 5H**). Similarly, patient 5 had CTCs with high EGFR expression, but-unlike other patients – this expression was significantly lower at the membrane (P < 0.0001, **Fig. 5I**). While these CTCs have high EGFR signalling that might otherwise indicate for monoclonal antibody inhibitors of EGFR, the lack of cell-permeability of these inhibitors means that they would not access intracellular EGFR and thus may prove ineffective for EGFR inhibition in this patient^74^.

### CTC subpopulation analysis highlights substantive intra-patient heterogeneity and possible personalised therapeutic strategies

Subcellular analysis of marker levels and localization demonstrates a distinguishable profile for each patient, yet intra-patient heterogeneity in CTC (sub)populations is also prominent. To identify and characterize patient-specific CTC subpopulations, UMAP embeddings were generated for CTCs from each patient followed by Hierarchical Density-Based Clustering of Applications with Noise (HDBSCAN)-mediated unsupervised cluster estimation (**Fig. 6A-D**; patient 2 presented only one cluster and is not analysed further in terms of subpopulations). For each patient, per CTC signalling pathway category profiles were radar plotted along with mean values (bold) per clustered CTC subpopulation (**Fig. 6A-D**; signalling pathways categories detailed in **Supplementary Table 3**), thereby delineating differences between CTC subpopulations. Random forest feature selection was performed to identify the top performing markers for distinguishing subpopulations within each patient’s CTC population, the top 10 are displayed in **Supplementary Fig. 6**. CTCs from patient 3 displayed relatively consistent pathway activations across clusters, with the exception of Cluster 3 (green) which showed increased active β-catenin signalling and increased markers of neuroendocrine differentiation (known to be driven by β-catenin^75^) (**Fig. 6A**). CTCs from patient 4 comprised 3 clusters, with Clusters 1 and 2 showing high TGFβR pathway activation, while Cluster displayed a distinctive profile with increased total and nuclear AR signal (**Fig. 6B**). Patient 5’s CTC clusters showed subtly different signatures, with Cluster 3 having increased activation levels across a range of pathways, potentially implying a resistant subtype (**Fig. 6C**). Perhaps most strikingly, subpopulation analysis of patient 6 CTCs (**Fig. 6D**), despite only detecting 4 CTCs in total, confirmed two separate CTC subpopulations, as indicated in earlier analyses (**Fig. 4B**). These two clusters correlate with the CTC groups associated either with therapy ‘responder’ or ‘non-responder’ CTC phenotypes observed in CTC UMAP embeddings including cells from all patients. More specifically, patient 6 Cluster 1 ‘responder’ CTCs show heightened p-STAT3 activity as well as increased CD44, KRAS and mTOR expression, while Cluster 2 CTCs align with the ‘non-responder’ phenotype and show heightened PSMA levels as well as increased total and nuclear AR (**Fig. 6D**); markers of AR-driven therapy resistance. This strategy elucidates and characterizes intra-patient heterogeneity which may be useful for selection combinatorial therapy.

**Figure 6:**
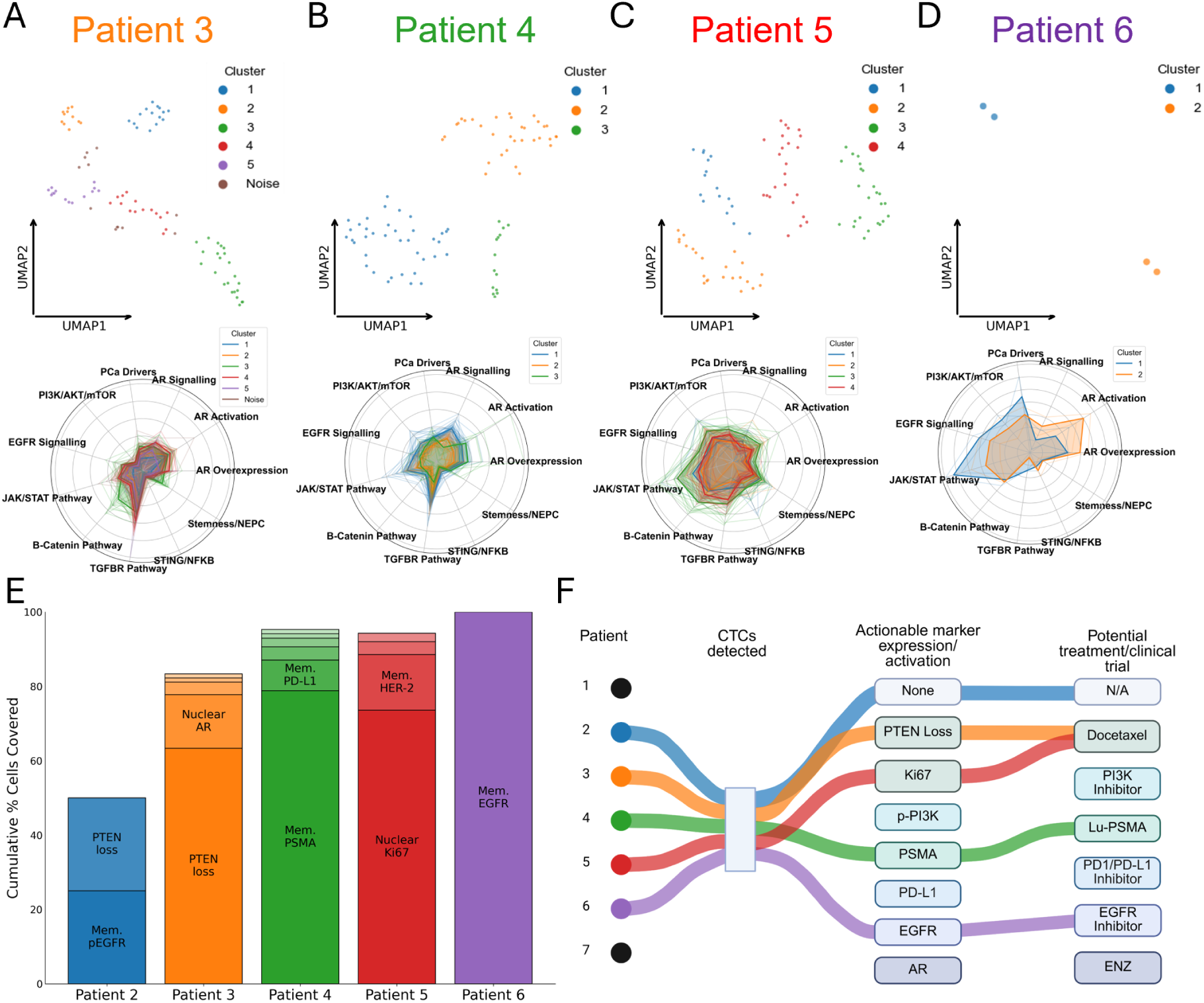
Patient CTC subpopulation analysis may guide personalised actionable treatment selection. (**A-D**) Per cell marker intensity UMAP-embedding with HDBSCAN clustering-defined CTC subpopulations from patients 3,4,5 and 6. Radar plots of signalling pathway category signatures for every CTC and the subpopulation mean (bold) of each cluster visualise differences in signalling profiles. (**E**) For each patient, a greedy analysis was performed on a reduced list of actionable biomarkers, iteratively selecting the biomarker with top-quartile expression in the most CTCs (bottom quartile for PTEN). (**F**) Potential treatments or clinical trial options guided by CTC actionable biomarker analyses per patient. Signalling pathway categories used in radar plots defined in **Supplementary Table 3**.

However, another approach is to instead identify which actionable signatures are conserved through all of a patients CTCs. To assess the potential utility of our deep multiplexing strategy for the future guidance of precision oncology, CTCs from each patient were interrogated using greedy analysis of a subset of established, actionable markers (**Fig. 6E**). This approach iteratively identified actionable markers with extreme expression (top or bottom quartile according to biomarker, i.e. targeting high or low expression) on the largest possible subset of CTCs within each patient, thus defining the largest targetable CTC subpopulation with a specific molecular vulnerability. We identified membrane EGFR, pEGFR, HER-2, PD-L1, PSMA or p-PI3K, and nuclear PTEN, Ki67 or AR, as well as cytoplasmic mTOR, as established biomarkers for predicting the efficacy of treatment approaches prevalent in PCa or other solid tumors^76–83^. This approach thus linked characterized biomarker expression to potential guidance of therapy selection (or clinical trial stratification) (**Fig. 6F**). For instance, CTCs from non-responder patient 3 displayed loss of PTEN and low tubulin expression; suggesting that taxane-based therapies could be beneficial^84^ (**Fig. 6E-F**). Patient 4 CTCs displayed high membrane PSMA, the direct binding target of Lutetium-177 PSMA (Lu-PSMA) therapy, indicating the potential efficacy of this targeted radio-ligand treatment^80^ (**Fig. 6E-F**). Conversely, responder patient 5 CTCs displayed virtually no surface PSMA, suggesting the unsuitability of Lu-PSMA, while high nuclear Ki67 suggests that instead chemotherapy may be beneficial^83^ (**Fig. E-F**). CTCs from therapy-responsive patients 5 and 6 exhibited high membrane EGFR expression and signaling, known correlates of EGFR inhibitor-effectiveness; potentially making these patients suitable candidates for EGFR-inhibitor trials in PCa^77^.

## Discussion

Precision oncology has transformative potential^85^ but requires reliable, repeatable diagnostics to identify and longitudinally monitor actionable molecular targets in evolving tumors^86,87^. Liquid biopsies like blood samples enable minimally invasive monitoring to guide anti-cancer treatment strategies. Among blood analytes, CTCs uniquely contain all tumor-derived molecular analytes^88^ while also capturing emergent molecular assembly into cellular-scale phenotypes^89,90^. CTCs are also functional biomarkers of metastasis itself^5,26^. However, CTC exploitation has been limited by low molecular marker plexity (4-5 markers/CTC) when using standard immunofluorescence methods, constraining accurate detection of heterogeneous CTCs as well as broad molecular profiling needed to guide the use of expanding therapeutic options. We have tackled this limitation through deep multiplexed imaging of (here) up to 50 molecular markers per CTC. This improves detection of CTCs, including those with non-canonical phenotypes, while also capturing the levels and functionally significant subcellular localizations of (phospho)protein markers for key phenotypic, oncogenic and resistance signals. This represents an order-of-magnitude advance beyond standard IF-based CTC profiling and provides proof-of-principle for a new end-to-end strategy that may enhance the potential of CTCs as molecular guides for precision oncology.

Our deep multiplexed imaging approach improves CTC identification via manual definition and via (subsequent) automated machine learning-based CTC classification. This is achieved by incorporating a range of markers that address the emerging diversity of CTCs^19–22^. Consequently, when considering marker expression levels-only, Boruta machine learning defined a top-ten ranking of markers providing maximal precision for CTC classification, especially when compared to canonical thresholding of DNA, CD45, EpCAM and Cytokeratin. Machine learning classification also leveraging marker subcellular localizations achieved almost the same performance whilst relying primarily on EpCAM and AR features, demonstrating how subcellular spatial resolution of marker localizations can contribute to efficient CTC detection. This emphasises the value of single-cell imaging over single-cell methods lacking subcellular spatial resolution. Notably, our strongest classification models emphasised inclusion of not just general cancer markers, but also markers with (here PCa) tissue selectivity/specificity, like AR and PSMA. For instance, including AR permitted identification of non-canonical CD45-positive CTCs that have been reported to evade immune responses and promote metastasis^22^, and are here defined in part by AR expression that is generally absent from normal blood cells^91^. We also note, for example, our detection of CTCs with neuroendocrine phenotypes, which might be missed using narrower CTC definitions. Taken together, the deep multiplexed CTC profiles presented here, including both expression levels and subcellular localizations, combined with machine learning to support higher precision and confidence in CTC detection while at the same time addressing more diverse CTC phenotypes that are reflective of the widening spectrum of CTC profiles reported in literature.

While our CTC identification here was predominantly guided by traditional CTC markers (with manual confirmation), future studies of larger CTC populations from increased patient numbers, including longitudinal sampling, will allow thorough testing of multivariate inclusion/exclusion criteria for marker signatures guiding optimised, diversified CTC detection. Importantly, where similar marker panels are used but understanding of CTC definitions evolve, our computationally-based CTC definitions are retrospectively tuneable, such that the latest insights can be back-applied to existing datasets. Automatable via machine learning, such criteria are vital to the translation of CTC analyses into diagnostic settings.

Beyond enhancing CTC detection, deep multiplexed imaging here provides scope to dramatically expand the sampling of known and putative biomarkers spanning alternate oncogenic/resistance signals and phenotypic states that may, in any given patient, help to define actionable strategies for treatment. CRPC exemplifies the need for such a diverse precision diagnostic capacity, since late-stage therapy-resistant PCa can be promoted or protected by multiple alternate driver/resistance pathways^35,36^ and cancer cell phenotypes^92^. Significantly, unlike mutational profiling or transcriptomic analyses^89^, the levels and subcellular localizations of (phospho)proteins assessed herein constitute direct (as opposed to indirect or inferred) biomarkers of actual signalling pathway activity and phenotypic states. Moreover, these functional biomarkers are in many cases themselves the explicit targets of therapeutic interventions (e.g. AR, pEGFR, PD-L1 etc).

As exemplified during CTC classification, the resolution of biomarker subcellular localizations again provides critical information to understand the detailed functional status of disease drivers and thereby guide enhanced precision oncology. For instance, we detected patients with heightened surface expression of pEGFR (patient 6) or PSMA (patient 4), suggesting suitability for EGFR-inhibitor-or Lutetium-PSMA-treatments, respectively. Likewise, the total mean intensity of actionable biomarkers did not always align with the proportion of targetable, membrane-localised protein, as highlighted with patient 3 having high total PSMA but low membrane availability. While rarely recorded in existing diagnostics reliant on mutational, transcriptional or total (phospho)protein-level data, the surface-presence (or absence) of these biomarkers is likely to be a major determinant of their vulnerability to therapies that bind their targets when surface-resident. Nuclear localization of AR in a subset of CTCs from patient 3 implies residual AR pathway activity that may contribute to a lack therapy response, while patient 6 displayed heightened expression of cytoplasmic β-catenin without increased nuclear β-catenin. This implies limited Wnt signalling activity, with this conclusion critically dependent on the discernment of cytoplasmic versus nuclear β-catenin localization.

Considering multivariate quantitative profiles combining protein levels, phosphorylation status and subcellular localizations across multiple biomarkers, as well as the intra-patient heterogeneity of CTC populations, an ever more comprehensive picture of each patient in this proof-of-principle cohort emerged. With all patients receiving standard-of-care combinations of AR inhibitors (ADT and enzalutamide), heightened AR signalling and expression was expected to be the clearest indicator of resistance; being reported in ∼80% of CRPC patients^76^. Accordingly, CTCs from therapy non-responsive patients 3 and 4 displayed higher AR expression, higher AR activation (AR nuclear localisation) and higher PSMA levels (a downstream product of AR signalling). This profile correlates with reports that PSMA expression increases with the emergence of mCRPC^93^. Conversely, therapy responsive patients 5 and 6 exhibited high EGFR/PI3K/AKT expression and phospho-signalling as well as exaggerated CTC population heterogeneity (high entropy). In these patients, ADT appeared effective in inhibiting AR signalling, but this selective pressure may have triggered a range of compensatory signals producing a spectrum of resistant CTC sub-populations^94^. Of note, patient 2 presented a mixed picture, with reduce blood PSA post-treatment implying therapy-response. Yet blood PSA levels in fact remained 100-fold higher than other therapy-responsive patients, while molecular profiles as well as CTC population entropy mimicked the reduced diversity observed in non-responsive patients. While requiring validation through a larger patient population, this result may reveal sensitivity to disease states in transition between therapy-response and therapy-resistance.

As detailed above, the quantitative analysis of intra-patient CTC population entropy reveals crucial insights into tumor cell heterogeneity; a major cause of therapy-failure^95,96^. Moreover, given longitudinal liquid biopsy, deep multiplexed CTC imaging may be useful for tracking heterogeneity-dynamics reflecting the interplay of treatment, selection and resistance. In addition to molecular-scale biomarker information, this analysis also reveals cellular scale phenotype and cellular fate, with this ‘dual’ capacity a key advantage over other liquid biopsy approaches, such as ctDNA analysis. Beyond this generalised analysis of CTC variation, detection of intra-patient CTC subpopulations with different signalling, resistance and/or phenotypic profiles has potential to guide personalised selection of combined therapies^97^, which are increasingly seen as more effective that monotherapies^98^. As exemplified in our greedy analysis approach (**Fig. 6E**), 4 out of 5 patients with sufficient CTCs (to assess multiple subpopulations) contained CTC subpopulations with biomarkers implying different actionable therapeutic vulnerabilities. This analysis was performed even with CTC numbers as low as 4 (patient 6, where two subpopulations were discernible), suggesting the potential for subpopulation analyses in patient samples with low CTC counts.

Importantly, our deep multiplexed analysis of patient-derived CTCs followed validation that this approach detected known oncogenic/resistance/phenotypic changes spanning a three-stage cell line model of PCa therapy-resistance that emulates longitudinal CTC-sampling across successive disease and treatment stages^37^. We detected known marker expression and phospho-signalling changes associated with castrate– and enzalutamide-resistance in this model, including increased AR, NFκB/STING and JAK/STAT^47,61^. Pseudotime analysis based on per cell marker levels-only highlighted continuum changes linked to resistance phenotypes, emphasising how combining longitudinal sampling and pseudotemporal analysis of intra-sample heterogeneity can build a picture of progressively emerging resistance. Incorporating subcellular localisation features into pseudotemporal analysis enhanced this picture further by, for example, showing how functionally significant localization changes (e.g. increased nuclear AR or plasma membrane PD-L1) emerge at the most extreme pseudotemporal ranges of enzalutamide-resistance. Likewise, including subcellular localization features into machine learning models for resistance-state classification also increased model accuracy and confidence, when considering either individual biomarkers or multi-molecular biomarker signatures (**Fig. 1E**). Once again, this reflects the critical role of subcellular localization in the molecular regulation of (bio)marker functions, and therefore the value of combining deep multiplexed marker detection with imaging-based subcellular marker localization^99,100^.

## Conclusion

Safely accessible via minimally invasive blood sampling, liquid biopsy-derived CTCs provide safe, longitudinal access to functional cancer biomarkers spanning molecular to cellular scales, achieving unique scope to guide precision oncology. Yet given their rarity, maximising data acquisition from each and every CTC is vital, not only to detect actionable biomarkers, but also to detect variation in biomarker status between and within individual patients. To exploit the unique precision diagnostic opportunity embodied in CTCs, we have presented proof-of-principle for the utility of a novel pipeline enabling deep multiplexed immunofluorescence imaging of CTCs. Using quantitative single-cell and (sub)population analyses, supported by machine learning-based classification and (biomarker) feature selection strategies, we have parsed molecular-scale oncogenic and resistance pathway signals, as well as cellular-scale phenotypes, to delineate examples of actionable biomarkers and biomarker signatures suggesting specific therapeutic choices. With follow-up and expansion of this and other cancer cohorts ongoing, we believe this approach presents one avenue for further exploration of the precision diagnostic potential of CTCs to support advances in precision oncology.

## Acknowledgements

I.G. and F.V.K. are supported by Australian Government Research Training Program (RTP) Scholarships. M.D. is supported by the UNSW University Postgraduate Award. F.V.K. receives a Top-Up award from SPHERE Cancer CAG and Cancer Institute NSW. D.P.N. is supported by a Tour de Cure Pioneering Grant (RSP-573-2024). J.G.L. is supported by a University of New South Wales Scientia Research Fellowship, a Ramaciotti Biomedical Research Award, an ARC Development Project grant (DP170103599). J.G.L, T.M.B, and T.L.R. are supported by NHMRC Ideas Grants (GNT1184009, GNT2012848, GNT2028506). J.G.L is supported by and the National Breast Cancer Foundation Collaborative Accelerator Grant (2024/CRA0020) a Tour de Cure Pioneering Grant (RSP-547-FY2023). We thank Rob Mann for their assistance editing the manuscript. We acknowledge the Katharina Gaus Light Microscopy Facility (KGLMF), part of the Mark Wainwright Analytical Centre at UNSW, for providing access to advanced light microscopy resources.

## Author contributions

Concept and design: J.G.L., T.L.R., T.M.B. and T.J.M.; development of methodology: T.J.M., Y.Z., T.K. and Y.M.; acquisition of data (sample processing, imaging): T.J.M., Y.Z., T.K., U.N. and A.J.; recruitment of patients: U.N., W.C. and P.dS.; analysis and interpretation of data (e.g. Image processing, machine learning and statistical analysis): T.J.M., T.H., D.N., F.K., I.G., F.V. and J.G.L.; Writing of manuscript: T.J.M., J.G.L. and T.M.B.; review and/or edit of manuscript: Y.M., F.K., I.H., P.dS. and T.L.R.

## Code availability

Code used in this manuscript is made available at https://github.com/CancerSystemsMicroscopyLab/CTC_Multiplexed_Immunofluorescence.g it

## Supplementary Tables

**Supplementary Table 1:** Granularity, Texture, Intensity and Ratio features captured for each cell segmentation of full cell, cytoplasm and nuclei.

| Granularity | Texture | Intensity | Ratio |
| --- | --- | --- | --- |
| Granularity_1 | Texture_AngularSecondMoment_3_00_256 | Intensity_IntegratedIntensity | Nuclear: Cytoplasm<br>Mean Intensity Ratio |
| Granularity_2 | Texture_AngularSecondMoment_3_01_256 | Intensity_IntegratedIntensityEdge | Nuclear: Cytoplasm<br>Integrated Intensity Ratio |
| Granularity_3 | Texture_AngularSecondMoment_3_02_256 | Intensity_LowerQuartileIntensity |  |
| Granularity_4 | Texture_AngularSecondMoment_3_03_256 | Intensity_MADIntensity |  |
| Granularity_5 | Texture_Contrast_3_00_256 | Intensity_MassDisplacement |  |
| Granularity_6 | Texture_Contrast_3_01_256 | Intensity_MaxIntensity |  |
| Granularity_7 | Texture_Contrast_3_02_256 | Intensity_MaxIntensityEdge |  |
| Granularity_8 | Texture_Contrast_3_03_256 | Intensity_MeanIntensity |  |
| Granularity_9 | Texture_Correlation_3_00_256 | Intensity_MeanIntensityEdge |  |
| Granularity_10 | Texture_Correlation_3_01_256 | Intensity_MedianIntensity |  |
| Granularity_11 | Texture_Correlation_3_02_256 | Intensity_MinIntensity |  |
| Granularity_12 | Texture_Correlation_3_03_256 | Intensity_MinIntensityEdge |  |
| Granularity_13 | Texture_DifferenceEntropy_3_00_256 | Intensity_StdIntensity |  |
| Granularity_14 | Texture_DifferenceEntropy_3_01_256 | Intensity_StdIntensityEdge |  |
| Granularity_15 | Texture_DifferenceEntropy_3_02_256 | Intensity_UpperQuartileIntensity |  |
| Granularity_16 | Texture_DifferenceEntropy_3_03_256 |  |  |
|  | Texture_DifferenceVariance_3_00_256 |  |  |
|  | Texture_DifferenceVariance_3_01_256 |  |  |
|  | Texture_DifferenceVariance_3_02_256 |  |  |
|  | Texture_DifferenceVariance_3_03_256 |  |  |
|  | Texture_Entropy_3_00_256 |  |  |
|  | Texture_Entropy_3_01_256 |  |  |
|  | Texture_Entropy_3_02_256 |  |  |
|  | Texture_Entropy_3_03_256 |  |  |
|  | Texture_InfoMeas1_3_00_256 |  |  |
|  | Texture_InfoMeas1_3_01_256 |  |  |
|  | Texture_InfoMeas1_3_02_256 |  |  |
|  | Texture_InfoMeas1_3_03_256 |  |  |
|  | Texture_InfoMeas2_3_00_256 |  |  |
|  | Texture_InfoMeas2_3_01_256 |  |  |
|  | Texture_InfoMeas2_3_02_256 |  |  |
|  | Texture_InfoMeas2_3_03_256 |  |  |
|  | Texture_InverseDifferenceMoment_3_00_256 |  |  |
|  | Texture_InverseDifferenceMoment_3_01_256 |  |  |
|  | Texture_InverseDifferenceMoment_3_02_256 |  |  |
|  | Texture_InverseDifferenceMoment_3_03_256 |  |  |
|  | Texture_SumAverage_3_00_256 |  |  |
|  | Texture_SumAverage_3_01_256 |  |  |
|  | Texture_SumAverage_3_02_256 |  |  |
|  | Texture_SumAverage_3_03_256 |  |  |
|  | Texture_SumEntropy_3_00_256 |  |  |
|  | Texture_SumEntropy_3_01_256 |  |  |
|  | Texture_SumEntropy_3_02_256 |  |  |
|  | Texture_SumEntropy_3_03_256 |  |  |
|  | Texture_SumVariance_3_00_256 |  |  |
|  | Texture_SumVariance_3_01_256 |  |  |
|  | Texture_SumVariance_3_02_256 |  |  |
|  | Texture_SumVariance_3_03_256 |  |  |
|  | Texture_Variance_3_00_256 |  |  |
|  | Texture_Variance_3_01_256 |  |  |
|  | Texture_Variance_3_02_256 |  |  |
|  | Texture_Variance_3_03_256 |  |  |

**Supplementary Table 2:** 86 Boruta features selected in Fig. 1C feature selection.

| Number | Feature |
| --- | --- |
| 1 | CellBody_Intensity_MeanIntensity_PSA |
| 2 | CellBody_Intensity_MeanIntensity_STING |
| 3 | CellBody_Intensity_MedianIntensity_PSA |
| 4 | CellBody_Intensity_UpperQuartileIntensity_PSA |
| 5 | CellBody_Intensity_UpperQuartileIntensity_STING |
| 6 | CellBody_Texture_Entropy_STING_3_00_256 |
| 7 | CellBody_Texture_Entropy_STING_3_01_256 |
| 8 | CellBody_Texture_Entropy_STING_3_02_256 |
| 9 | CellBody_Texture_SumAverage_PSA_3_00_256 |
| 10 | CellBody_Texture_SumAverage_PSA_3_01_256 |
| 11 | CellBody_Texture_SumAverage_PSA_3_02_256 |
| 12 | CellBody_Texture_SumAverage_PSA_3_03_256 |
| 13 | CellBody_Texture_SumAverage_STING_3_00_256 |
| 14 | CellBody_Texture_SumAverage_STING_3_01_256 |
| 15 | CellBody_Texture_SumAverage_STING_3_02_256 |
| 16 | CellBody_Texture_SumAverage_STING_3_03_256 |
| 17 | CellBody_Texture_SumEntropy_STING_3_00_256 |
| 18 | CellBody_Texture_SumEntropy_STING_3_02_256 |
| 19 | Cytoplasm_Intensity_MaxIntensityEdge_Vimentin |
| 20 | Cytoplasm_Intensity_MeanIntensityEdge_STING |
| 21 | Cytoplasm_Intensity_MeanIntensityEdge_Vimentin |
| 22 | Cytoplasm_Intensity_MeanIntensity_STING |
| 23 | Cytoplasm_Intensity_UpperQuartileIntensity_STING |
| 24 | Cytoplasm_Intensity_UpperQuartileIntensity_Vimentin |
| 25 | Cytoplasm_Texture_SumAverage_STING_3_01_256 |
| 26 | Cytoplasm_Texture_SumAverage_STING_3_02_256 |
| 27 | Nuclei_Intensity_MaxIntensityEdge_Vimentin |
| 28 | Nuclei_Intensity_MaxIntensity_Vimentin |
| 29 | Nuclei_Intensity_MeanIntensityEdge_Vimentin |
| 30 | Nuclei_Intensity_MeanIntensity_PSA |
| 31 | Nuclei_Intensity_MeanIntensity_Vimentin |
| 32 | Nuclei_Intensity_MedianIntensity_PSA |
| 33 | Nuclei_Intensity_MinIntensityEdge_DAPI |
| 34 | Nuclei_Intensity_UpperQuartileIntensity_PSA |
| 35 | Nuclei_Intensity_UpperQuartileIntensity_Vimentin |
| 36 | Nuclei_Texture_SumAverage_PSA_3_00_256 |
| 37 | Nuclei_Texture_SumAverage_PSA_3_01_256 |
| 38 | Nuclei_Texture_SumAverage_PSA_3_02_256 |
| 39 | Nuclei_Texture_SumAverage_PSA_3_03_256 |
| 40 | CellBody_Intensity_MADIntensity_PSA |
| 41 | CellBody_Intensity_MaxIntensity_Vimentin |
| 42 | CellBody_Texture_Entropy_PSA_3_02_256 |
| 43 | Cytoplasm_Texture_SumAverage_STING_3_00_256 |
| 44 | Cytoplasm_Texture_SumAverage_STING_3_03_256 |
| 45 | Nuclei_Granularity_16_p-Jak |
| 46 | CellBody_Intensity_MeanIntensity_Vimentin |
| 47 | CellBody_Texture_Entropy_PSA_3_00_256 |
| 48 | CellBody_Texture_Entropy_PSA_3_01_256 |
| 49 | CellBody_Texture_Entropy_STING_3_03_256 |
| 50 | Cytoplasm_Intensity_MaxIntensity_Vimentin |
| 51 | Nuclei_Intensity_MedianIntensity_Vimentin |
| 52 | CellBody_Granularity_16_p-Jak |
| 53 | CellBody_Intensity_MeanIntensityEdge_HER-2 |
| 54 | CellBody_Texture_SumEntropy_STING_3_01_256 |
| 55 | Cytoplasm_Intensity_MeanIntensityEdge_HER-2 |
| 56 | Nuclei_Intensity_LowerQuartileIntensity_PSA |
| 57 | CellBody_Intensity_UpperQuartileIntensity_Vimentin |
| 58 | CellBody_Texture_SumEntropy_PSA_3_00_256 |
| 59 | Nuclei_Intensity_MADIntensity_PSA |
| 60 | Nuclei_Intensity_MADIntensity_Vimentin |
| 61 | CellBody_Texture_AngularSecondMoment_PSA_3_02_256 |
| 62 | CellBody_Texture_SumEntropy_PSA_3_02_256 |
| 63 | Nuclei_Intensity_UpperQuartileIntensity_STING |
| 64 | CellBody_Texture_AngularSecondMoment_PSA_3_00_256 |
| 65 | CellBody_Texture_SumEntropy_STING_3_03_256 |
| 66 | Cytoplasm_Intensity_MeanIntensity_Vimentin |
| 67 | Nuclei_Texture_SumAverage_STING_3_03_256 |
| 68 | CellBody_Intensity_MeanIntensityEdge_STING |
| 69 | Nuclei_Texture_SumAverage_STING_3_02_256 |
| 70 | CellBody_Intensity_MeanIntensityEdge_Vimentin |
| 71 | CellBody_Texture_AngularSecondMoment_PSA_3_01_256 |
| 72 | Nuclei_Texture_SumAverage_STING_3_01_256 |
| 73 | CellBody_Texture_Entropy_PSA_3_03_256 |
| 74 | Cytoplasm_Granularity_16_p-Jak |
| 75 | Nuclei_Texture_SumAverage_STING_3_00_256 |
| 76 | Cytoplasm_Intensity_MedianIntensity_B-catenin (total) |
| 77 | Nuclei_Intensity_StdIntensity_Vimentin |
| 78 | CellBody_Granularity_16_Vimentin |
| 79 | CellBody_Texture_DifferenceVariance_B-catenin (total) 3 03 256 |
| 80 | Nuclei_Granularity_16_Vimentin |
| 81 | Cytoplasm_Intensity_UpperQuartileIntensity_B-catenin (total) |
| 82 | Cytoplasm_Granularity_16_Vimentin |
| 83 | Cytoplasm_Intensity_MeanIntensity_B-catenin (total) |
| 84 | Cytoplasm_Intensity_StdIntensityEdge_Vimentin |
| 85 | Nuclei_Intensity_MinIntensity_DAPI |
| 86 | CellBody_Texture_AngularSecondMoment_PSA 3 03 256 |

**Supplementary Table 3:**
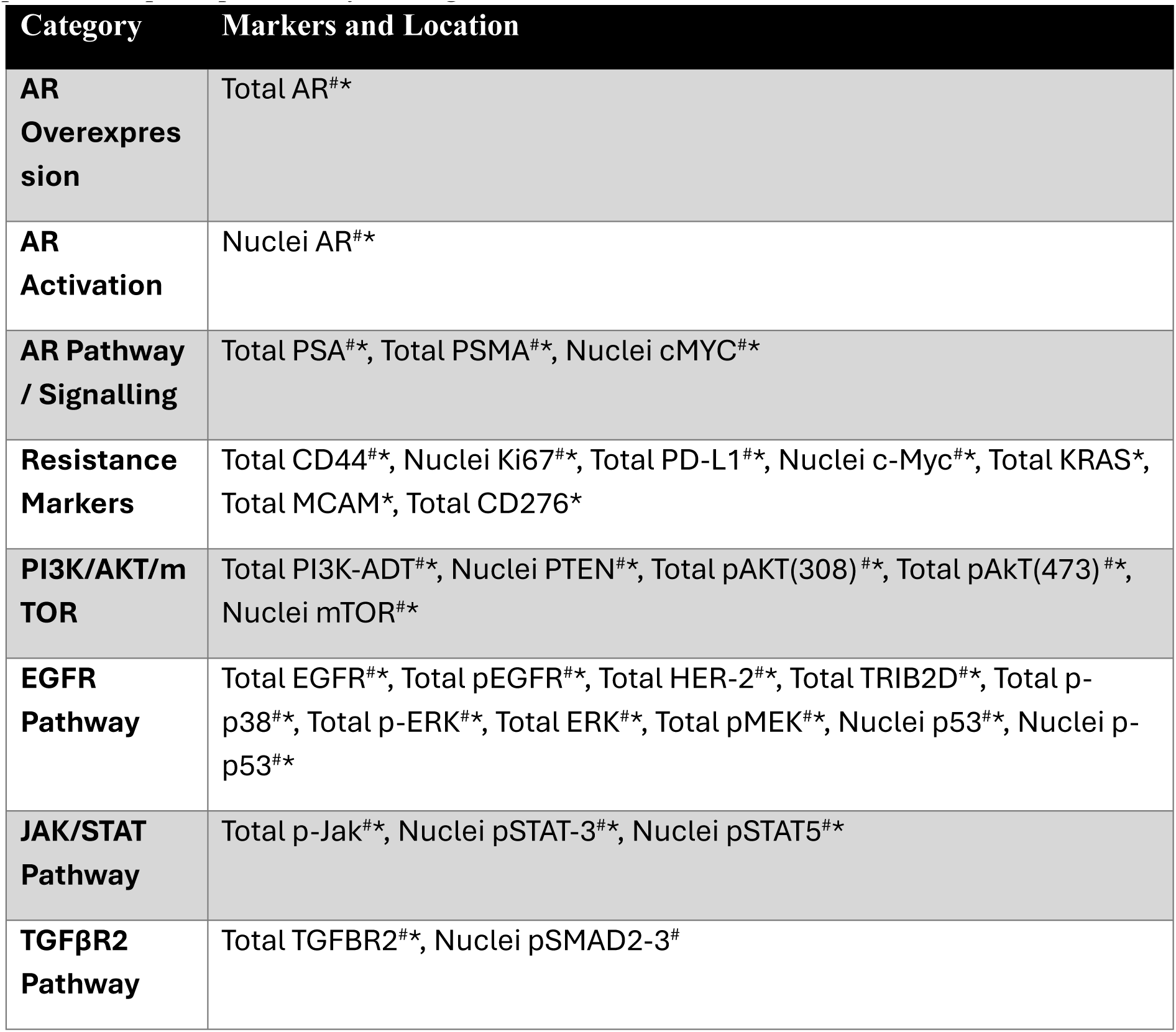
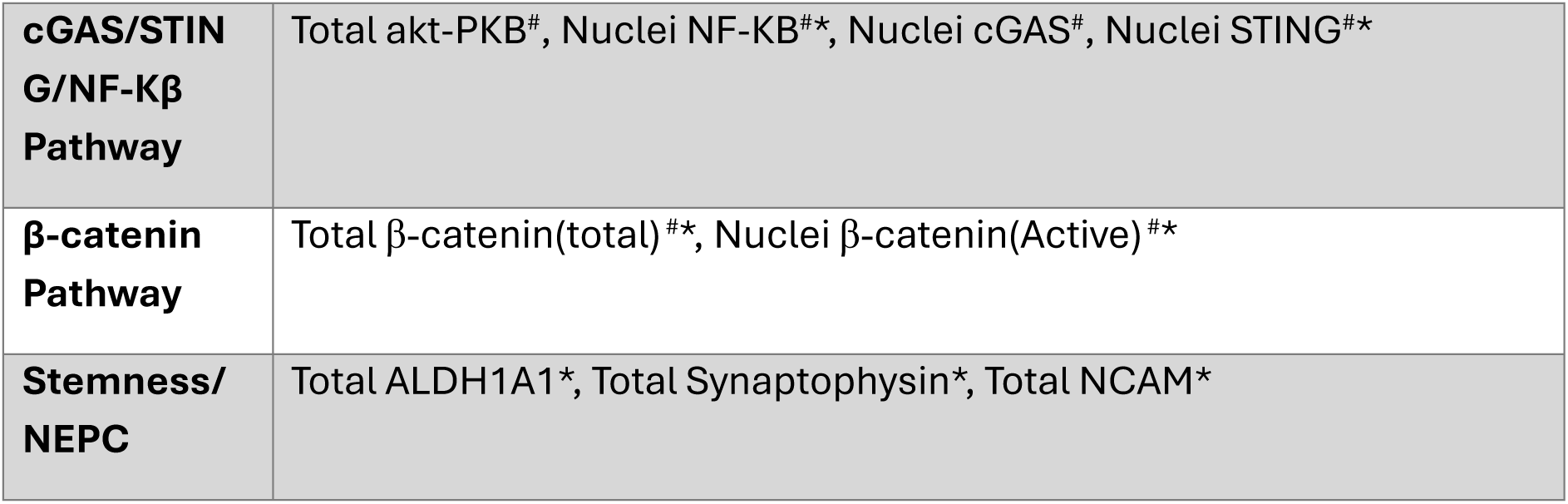
Total or nuclei mean marker intensity used for each pathway in radar plots. # denotes markers from cell line experiments from Fig. 2, * denotes markers for patient sample experiments from Fig. 3-4.

## Supplementary Figures

**Supplementary Figure 1:**
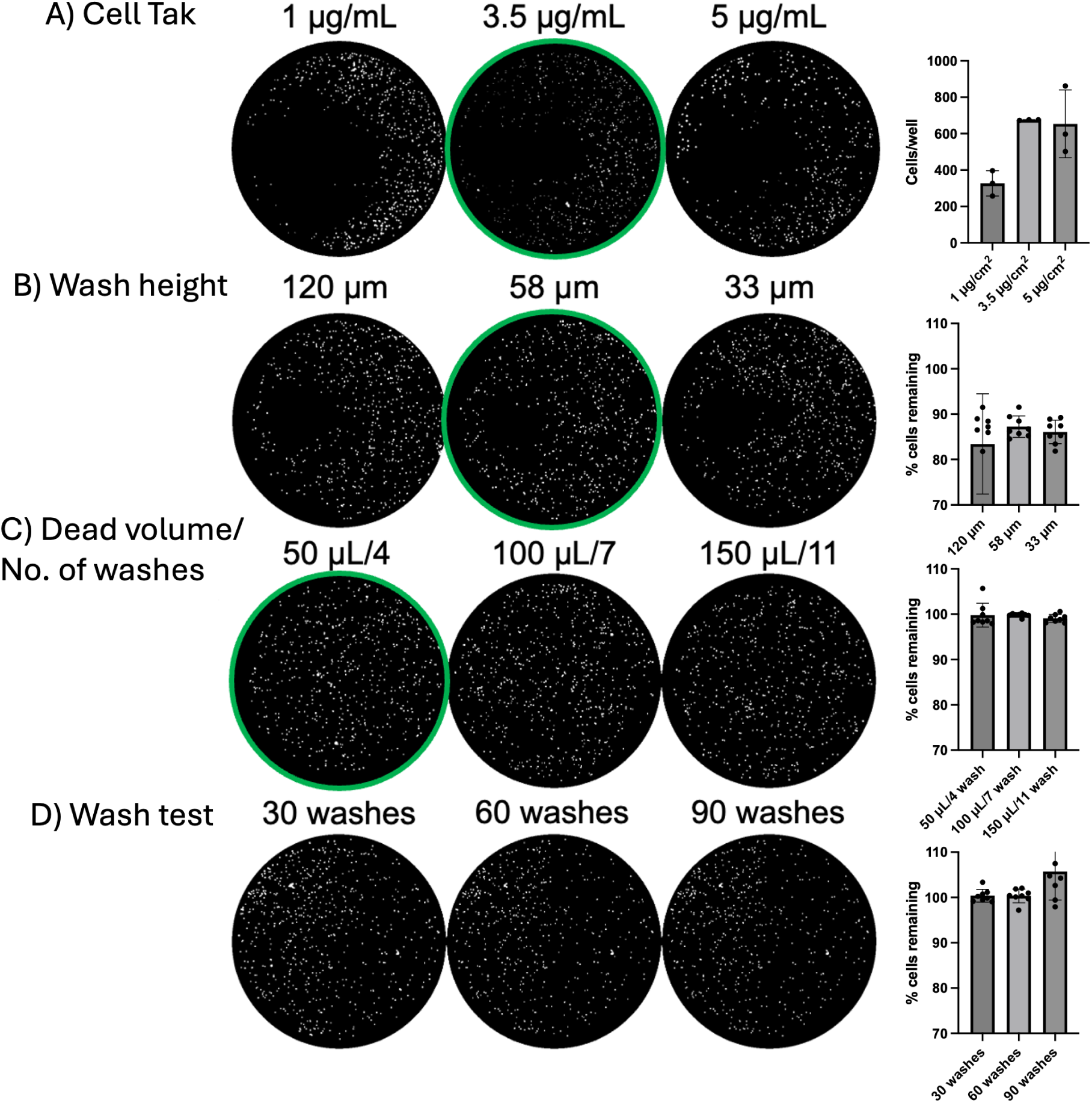
Optimization of sample washing. LNCaP cells were prepared in the same manner as CTC samples (fixed, frozen, thawed, washed) and seeded at 1000 cells/well. **A**) Cell Tak was tested at 1, 3.5 and 5 µg/cm^2^, with 3.5 µg/cm^2^ having the highest and most consistent retention of cells (n = 3). **B**) Using the 50 TS plate washer, the height of washing was tested at 120, 58 and 33 µm, with 58 µm having the highest retention of cells, without negatively affecting dead volume (n = 8). (**C**) After reducing the speed of washing, the dead volume of washing was tested at 50, 100 and 150 µL with 4, 7, and 11 washes respectively, to maintain similar dilution factors. 50 µL showed no significant decrease in cell number, while allowing for optimal washing (n = 8). (**D**) Under these conditions, no significant cell loss was observed after 30, 60 or 90 washes (n = 8).

**Supplementary Figure 2:**
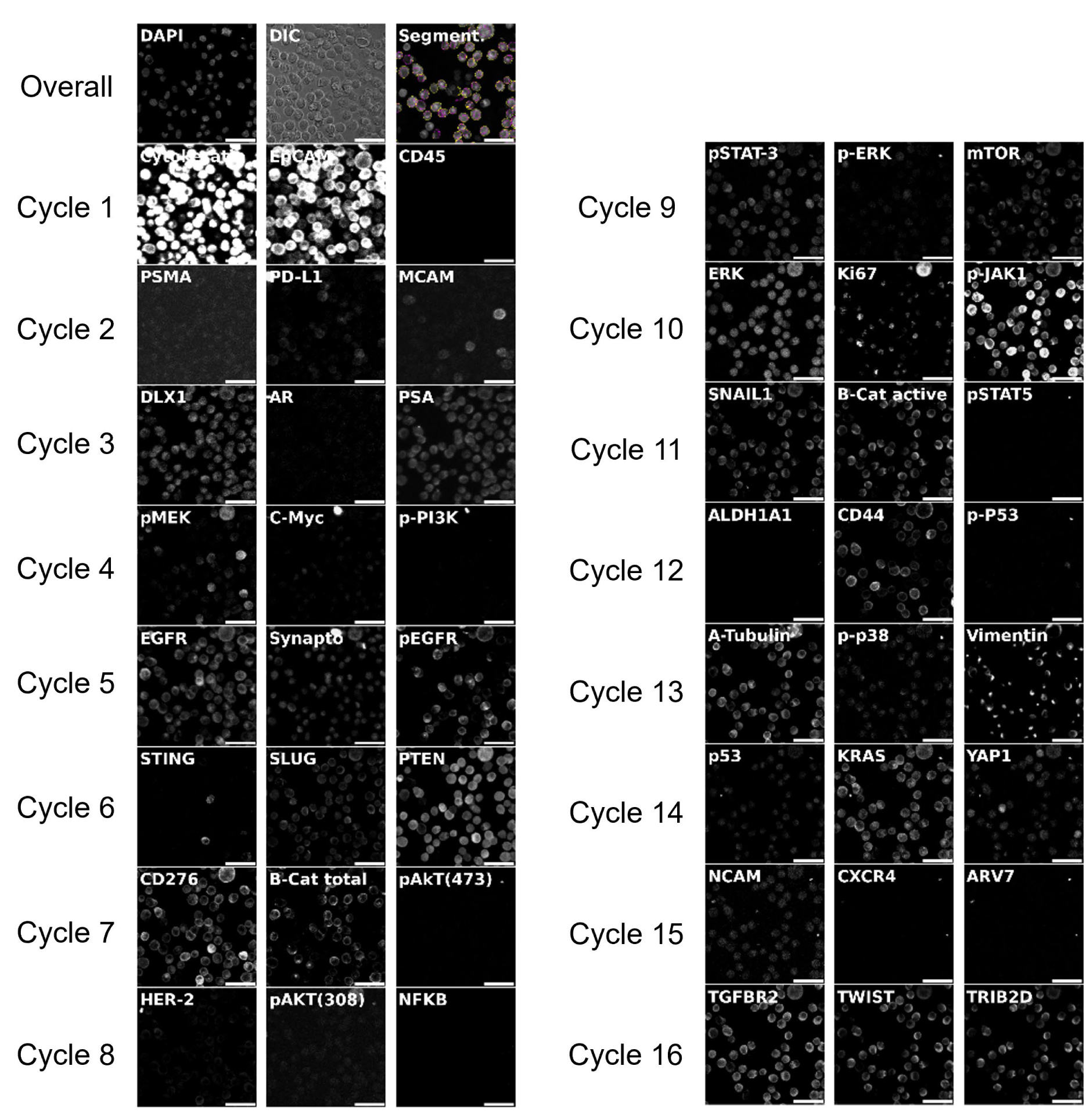
DU145 control cells labelled with antibodies used for CycIF in Figs 3-4. Cytokeratin and EpCAM were set to levels for CTC detection, resulting in overexposure in DU145.

**Supplementary Figure 3:**
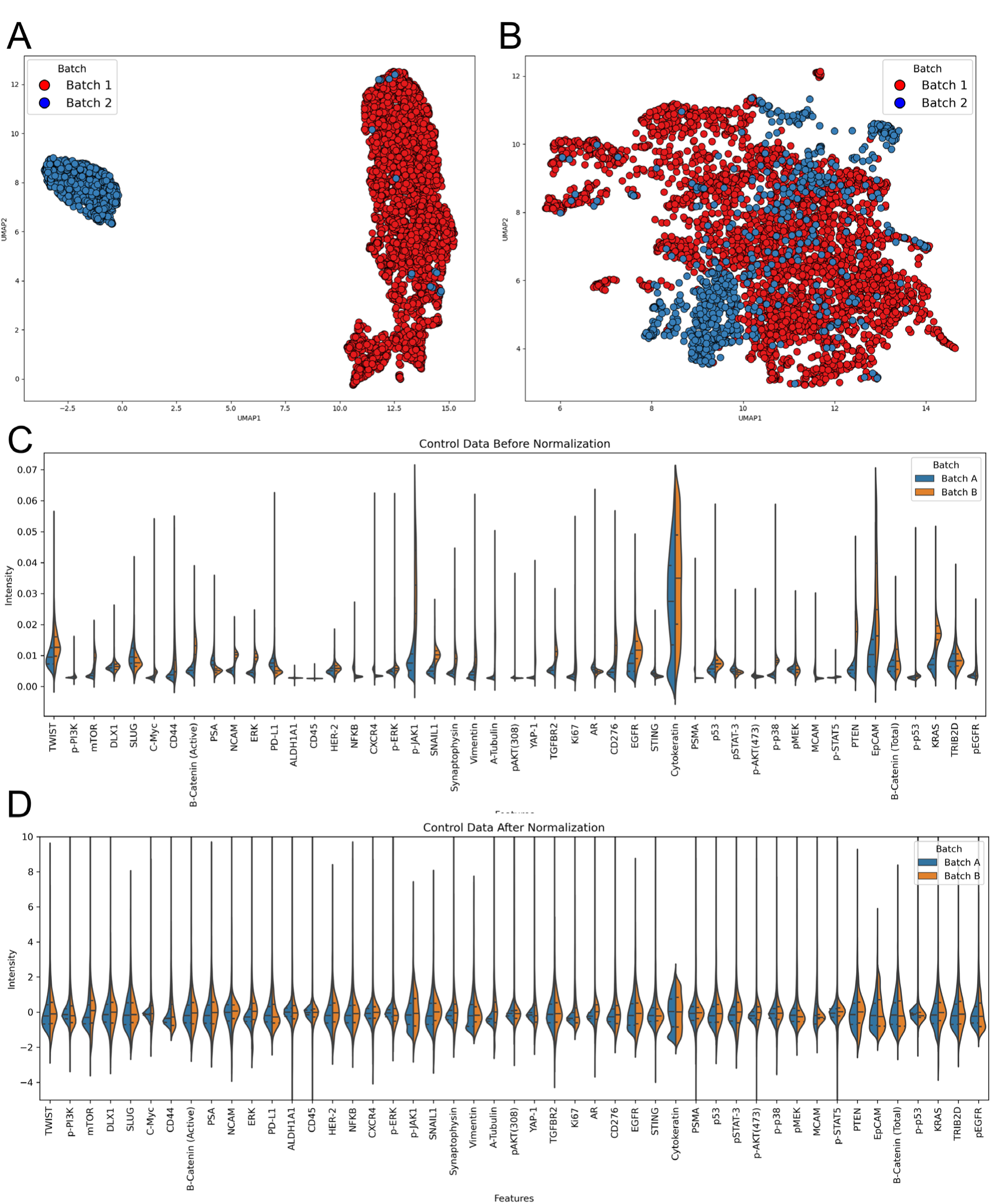
Batch normalisation. Aliquots of fixed, frozen DU145 PCa cells from a single passage were labelled and imaged in each of two CTC processing batches to allow for robust Z-score-based intensity normalization. UMAP of DU145 cells from each processing batch before (**A**) and after (**B**) normalization. Mean intensity distributions of markers in DU145 cells from each processing batch before (**C**) and after (**D**) normalization.

**Supplementary Figure 4:**
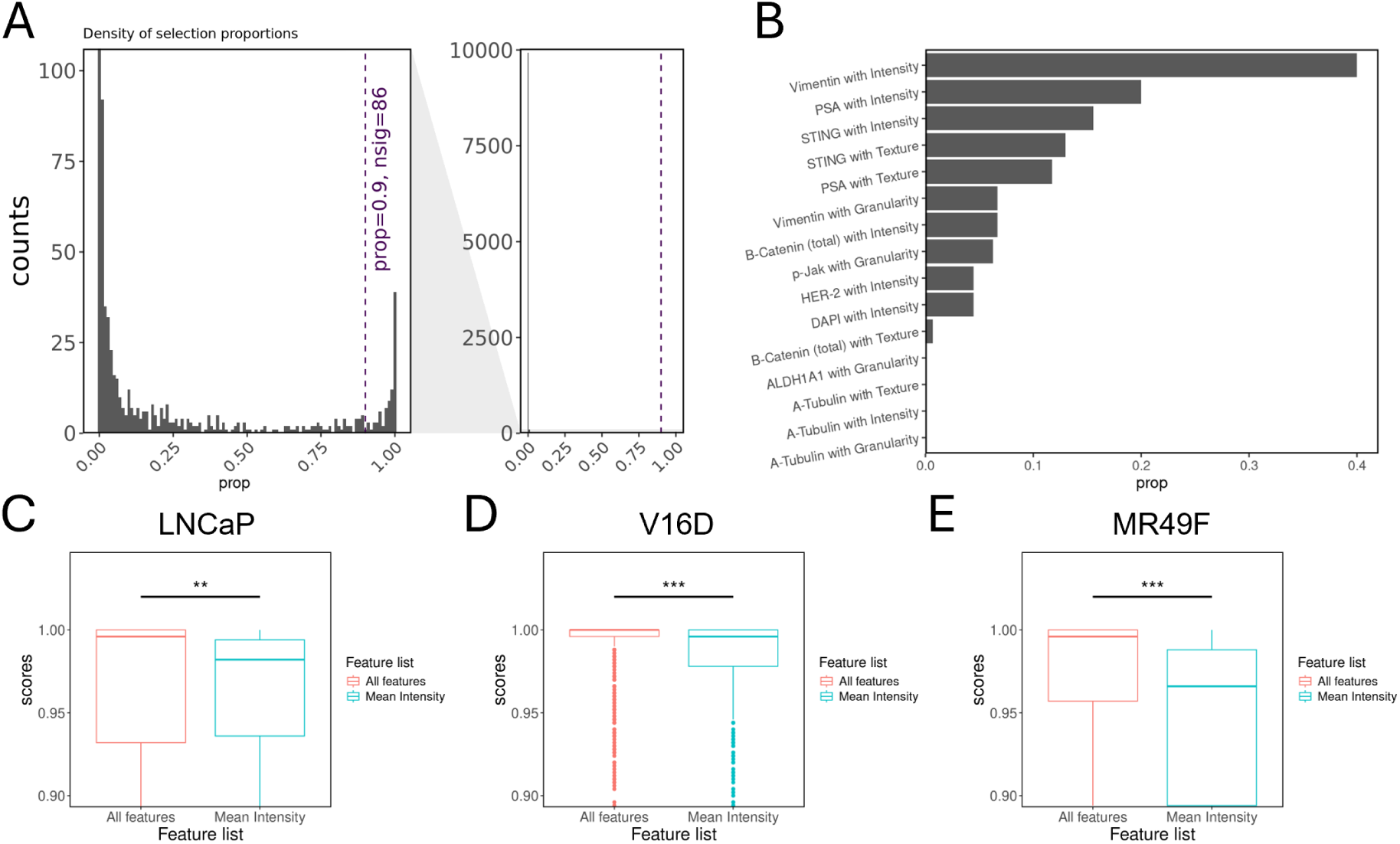
Boruta feature selection. (**A**) Histogram showing the proportion times each feature was selection via Boruta across all iterations, with 86 features selected above the 0.9 threshold. (**B**) Marker and feature group (Intensity, granularity or texture) that was selected via Boruta. (**C**) Comparison of prediction confidence from two multiclass random forest classifiers using with all features or mean intensity to predict LNCaP, (**D**) V16D or (**E**) MR49F (Mann-Whitney statistical test).

**Supplementary Figure 5:**
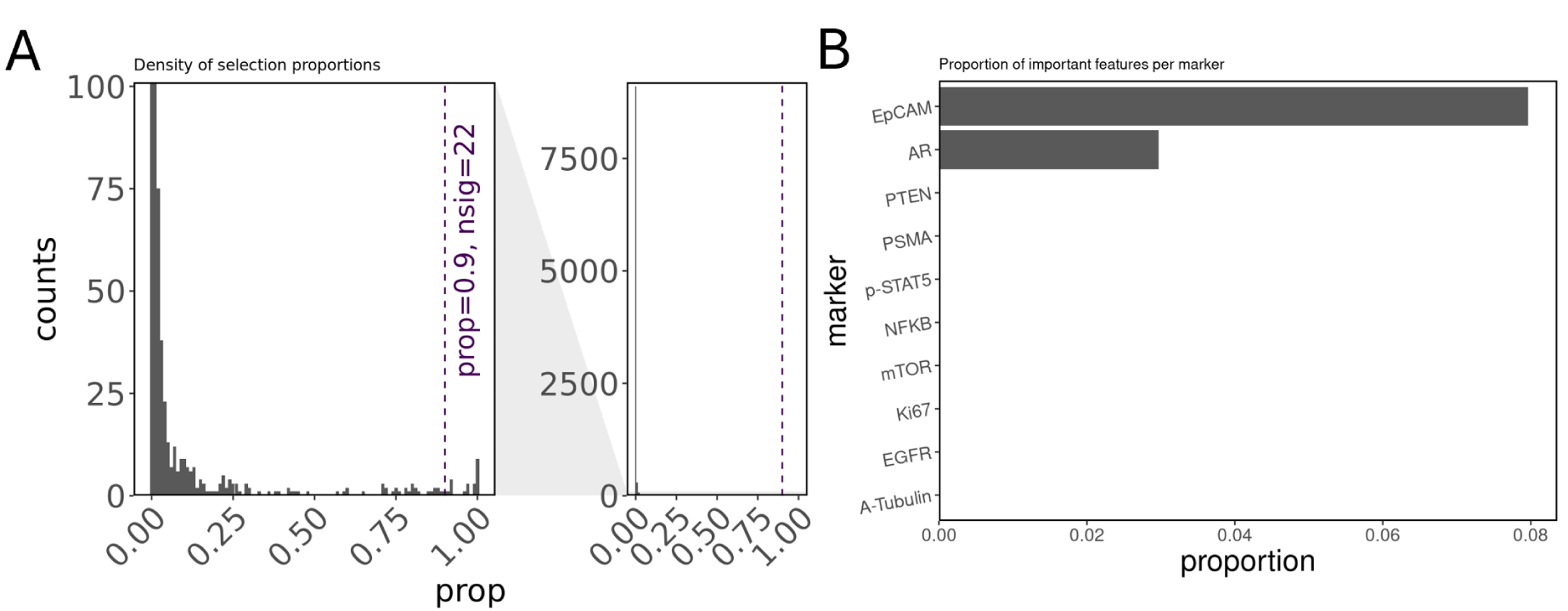
(**A**) Histogram showing the proportion times each feature was selection via Boruta for CTC identification across all iterations, with 22 features selected above the 0.9 threshold. (**B**) proportion of important feature per marker that was selected via Boruta.

**Supplementary Figure 6:**
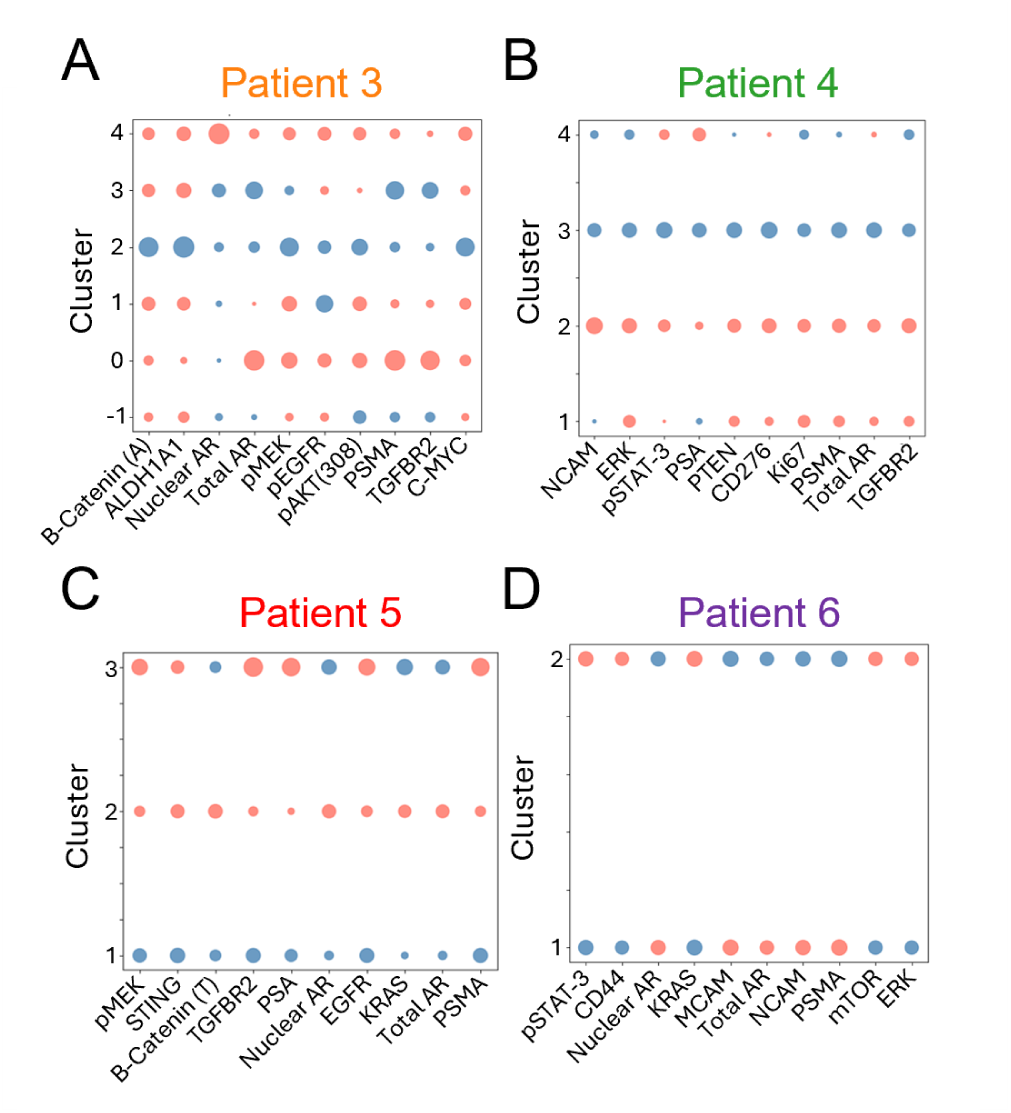
(**A-D**) The top 10 discriminative features of each sub-patient clusters from Fig 6 of patients 3,4,5 and 6, respectively.

## Notes

### Competing Interest Statement

The authors have declared no competing interest.

## References

1. Chae YK, PanAP, Davis AA et al. Path toward precision oncology: review of targeted therapy studies and tools to aid in defining ‘actionability’ of a molecular lesion and patient management support. Mol. Cancer Ther.16(12), 2645–2655 (2017).

2. Lassen UN, Makaroff LE, Stenzinger A, Italiano A, Vassal G, Garcia-Foncillas J, Avouac B. Precision oncology: a clinical and patient perspective. Future Oncology. 2021 Jun;17(30):3995–4009.

3. Liu, B., Zhou, H., Tan, L. et al. Exploring treatment options in cancer: tumor treatment strategies. Sig Transduct Target Ther 9, 175 (2024). 10.1038/s41392-024-01856-7

4. Wahida, A., Buschhorn, L., Fröhling, S. et al. The coming decade in precision oncology: six riddles. Nat Rev Cancer 23, 43–54 (2023).

5. Alix-Panabières, C., and Pantel, K. (2021). Liquid Biopsy: From Discovery to Clinical Application. Cancer Discov. 11, 858–873. 10.1158/2159-8290.CD-20-1311.

6. Gu, X., Wei, S. and Lv, X., 2024. Circulating tumor cells: from new biological insights to clinical practice. Signal transduction and targeted therapy, 9(1), p.226.

7. Payne, K., Brooks, J., Batis, N., Khan, N., El-Asrag, M., Nankivell, P., Mehanna, H. and Taylor, G., 2023. Feasibility of mass cytometry proteomic characterisation of circulating tumour cells in head and neck squamous cell carcinoma for deep phenotyping. British Journal of Cancer, 129(10), pp.1590–1598.

8. Scher, H.I., Graf, R.P., Schreiber, N.A., McLaughlin, B., Lu, D., Louw, J., Danila, D.C., Dugan, L., Johnson, A., Heller, G. and Fleisher, M., 2017. Nuclear-specific AR-V7 protein localization is necessary to guide treatment selection in metastatic castration-resistant prostate cancer. European urology, 71(6), pp.874–882.

9. Hofman, P., Heeke, S., Alix-Panabières, C. and Pantel, K., 2019. Liquid biopsy in the era of immuno-oncology: is it ready for prime-time use for cancer patients?. Annals of Oncology, 30(9), pp.1448–1459.

10. Gorges, T.M., Riethdorf, S., von Ahsen, O., Nastały, P., Röck, K., Boede, M., Peine, S., Kuske, A., Schmid, E., Kneip, C. and König, F., 2016. Heterogeneous PSMA expression on circulating tumor cells-a potential basis for stratification and monitoring of PSMA-directed therapies in prostate cancer. Oncotarget, 7(23), p.34930.

11. Fehm, T., Müller, V., Aktas, B., Janni, W., Schneeweiss, A., Stickeler, E., Lattrich, C., Löhberg, C.R., Solomayer, E., Rack, B. and Riethdorf, S., 2010. HER2 status of circulating tumor cells in patients with metastatic breast cancer: a prospective, multicenter trial. Breast cancer research and treatment, 124(2), pp.403–412.

12. Jordan, N.V., Bardia, A., Wittner, B.S., Benes, C., Ligorio, M., Zheng, Y., Yu, M., Sundaresan, T.K., Licausi, J.A., Desai, R. and O’Keefe, R.M., 2016. HER2 expression identifies dynamic functional states within circulating breast cancer cells. Nature, 537(7618), pp.102–106.

13. Babayan, A., Hannemann, J., Spoetter, J., Mueller, V., Pantel, K. and Joosse, S.A., 2013. Heterogeneity of estrogen receptor expression in circulating tumor cells from metastatic breast cancer patients. PloS one, 8(9), p.e75038.

14. Visal TH, den Hollander P, Cristofanilli M, Mani SA. Circulating tumour cells in the – omics era: how far are we from achieving the ‘singularity’? Br J Cancer. 2022 Jul;127(2):173–184. doi: 10.1038/s41416-022-01768-9. Epub 2022 Mar 10. PMID: 35273384; PMCID: PMC9296521.

15. Allen TA. The Role of Circulating Tumor Cells as a Liquid Biopsy for Cancer: Advances, Biology, Technical Challenges, and Clinical Relevance. Cancers (Basel). 2024 Mar 31;16(7):1377. doi: 10.3390/cancers16071377. PMID: 38611055; PMCID: PMC11010957.

16. Thiele, J.A., Bethel, K., Králíčková, M. and Kuhn, P., 2017. Circulating tumor cells: fluid surrogates of solid tumors. Annual Review of Pathology: Mechanisms of Disease, 12(1), pp.419–447.

17. Lawrence, R., Watters, M., Davies, C.R. et al. Circulating tumour cells for early detection of clinically relevant cancer. Nat Rev Clin Oncol 20, 487–500 (2023). 10.1038/s41571-023-00781-y

18. Lin, D., Shen, L., Luo, M. et al. Circulating tumor cells: biology and clinical significance. Sig Transduct Target Ther 6, 404 (2021). 10.1038/s41392-021-00817-8

19. Menyailo, M.E., Tretyakova, M.S. and Denisov, E.V., 2020. Heterogeneity of circulating tumor cells in breast cancer: identifying metastatic seeds. International Journal of Molecular Sciences, 21(5), p.1696.

20. Pecot, C., Bischoff, F., Mayer, J., Wong, K., Pham, T., Bottsford-Miller, J., Stone, R., Lin, Y., Jaladurgam, P., Roh, J., Goodman, B., Merritt, W., Pircher, T., Mikolajczyk, S., Nick, A., Celestino, J., Eng, C., Ellis, L., Deavers, M., & Sood, A. (2011). A novel platform for detection of CK+ and CK-CTCs.. Cancer discovery, 1 7, 580–6. 10.1158/2159-8290.CD-11-0215.

21. Reduzzi, C., Gerratana, L., Zhang, Y., D’Amico, P., Shah, A., Davis, A., Manai, M., Silvestri, M., Zhang, Q., Donahue, J., & Cristofanilli, M. (2022). CK+/CD45+ (dual-positive) circulating cells are associated with prognosis in patients with advanced breast cancer.. Journal of Clinical Oncology. 10.1200/jco.2022.40.16_suppl.1093.

22. Yang, C., Wang, X., To, K.K., Cui, C., Luo, M., Wu, S., Huang, L., Fu, K., Pan, C., Liu, Z. and Fan, T., 2024. Circulating tumor cells shielded with extracellular vesicle-derived CD45 evade T cell attack to enable metastasis. Signal Transduction and Targeted Therapy, 9(1), p.84

23. Cha, J., Cho, H., Chung, J., Park, J., & Han, K. (2023). Effective Circulating Tumor Cell Isolation Using Epithelial and Mesenchymal Markers in Prostate and Pancreatic Cancer Patients. Cancers, 15. 10.3390/cancers15102825.

24. Wu, S., Liu, S., Liu, Z., Huang, J., Pu, X., Li, J., Yang, D., Deng, H., Yang, N., & Xu, J. (2015). Classification of Circulating Tumor Cells by Epithelial-Mesenchymal Transition Markers. PLoS ONE, 10. 10.1371/journal.pone.0123976.

25. Zhao, X., Wang, Z., Chen, C., Di, L., Bi, Z., Li, Z., & Liu, Y. (2019). Molecular detection of epithelial-mesenchymal transition markers in circulating tumor cells from pancreatic cancer patients: Potential role in clinical practice. World Journal of Gastroenterology, 25, 138–150. 10.3748/wjg.v25.i1.138.

26. Zhang, X., Wei, L., Li, J., Zheng, J., Zhang, S. and Zhou, J., 2019. Epithelial mesenchymal transition phenotype of circulating tumor cells is associated with distant metastasis in patients with NSCLC. Molecular Medicine Reports, 19(1), pp.601–608.

27. Bastos, D., & Antonarakis, E. (2018). CTC-derived AR-V7 detection as a prognostic and predictive biomarker in advanced prostate cancer. Expert Review of Molecular Diagnostics, 18, 155–163. 10.1080/14737159.2018.1427068.

28. Scher, H., Lu, D., Schreiber, N., Louw, J., Graf, R., Vargas, H., Johnson, A., Jendrisak, A., Bambury, R., Danila, D., McLaughlin, B., Wahl, J., Greene, S., Heller, G., Marrinucci, D., Fleisher, M., & Dittamore, R. (2016). Association of AR-V7 on Circulating Tumor Cells as a Treatment-Specific Biomarker With Outcomes and Survival in Castration-Resistant Prostate Cancer. JAMA oncology, 2 11, 1441–1449. 10.1001/jamaoncol.2016.1828.

29. Mazel, M., Jacot, W., Pantel, K., Bartkowiak, K., Topart, D., Cayrefourcq, L., Rossille, D., Maudelonde, T., Fest, T., & Alix-Panabières, C. (2015). Frequent expression of PD-L1 on circulating breast cancer cells. Molecular Oncology, 9. 10.1016/j.molonc.2015.05.009.

30. Spiliotaki, M., Neophytou, C., Vogazianos, P., Stylianou, I., Gregoriou, G., Constantinou, A., Deltas, C., & Charalambous, H. (2022). Dynamic monitoring of PD-L1 and Ki67 in circulating tumor cells of metastatic non-small cell lung cancer patients treated with pembrolizumab. Molecular Oncology, 17, 792–809. 10.1002/1878-0261.13317.

31. Ilié, M., Szafer-Glusman, E., Hofman, V., Chamorey, E., Lalvée, S., Selva, E., Leroy, S., Marquette, C., Kowanetz, M., Hedge, P., Punnoose, E., & Hofman, P. (2018). Detection of PD-L1 in circulating tumor cells and white blood cells from patients with advanced non-small-cell lung cancer. Annals of Oncology, 29, 193–199. 10.1093/annonc/mdx636.

32. Lin, J.-R., Fallahi-Sichani, M. & Sorger, P. K. Highly multiplexed imaging of single cells using a high-throughput cyclic immunofluorescence method. Nat. Commun. 6, ncomms9390 (2015).

33. Xu, L., Mao, X., Guo, T., Chan, P.Y., Shaw, G., Hines, J., Stankiewicz, E., Wang, Y., Oliver, R.T.D., Ahmad, A.S. and Berney, D., 2017. The novel association of circulating tumor cells and circulating megakaryocytes with prostate cancer prognosis. Clinical Cancer Research, 23(17), pp.5112–5122.

34. Kaldjian, E.P., Ramirez, A.B., Sun, Y., Campton, D.E., Werbin, J.L., Varshavskaya, P., Quarre, S., George, T., Madan, A., Blau, C.A. and Seubert, R., 2018. The RareCyte® platform for next-generation analysis of circulating tumor cells. Cytometry Part A, 93(12), pp.1220–1225.

35. Wadosky KM, Koochekpour S. Molecular mechanisms underlying resistance to androgen deprivation therapy in prostate cancer. Oncotarget. 2016 Sep 27;7(39):64447–64470. doi: 10.18632/oncotarget.10901. PMID: 27487144; PMCID: PMC5325456.

36. Katzenwadel A, Wolf P. Androgen deprivation of prostate cancer: Leading to a therapeutic dead end. Cancer letters. 2015 Oct 10;367(1):12–7.

37. Kuruma, H., Matsumoto, H., Shiota, M., Bishop, J., Lamoureux, F., Thomas, C., Briere, D., Los, G., Gleave, M., Fanjul, A. and Zoubeidi, A., 2013. A novel antiandrogen, Compound 30, suppresses castration-resistant and MDV3100-resistant prostate cancer growth in vitro and in vivo. Molecular cancer therapeutics, 12(5), pp.567–576.

38. Quintanal-Villalonga, Á., Chan, J.M., Yu, H.A., Pe’er, D., Sawyers, C.L., Sen, T. and Rudin, C.M., 2020. Lineage plasticity in cancer: a shared pathway of therapeutic resistance. Nature reviews Clinical oncology, 17(6), pp.360–371.

39. Berthold, M.R., Cebron, N., Dill, F., Gabriel, T.R., Kötter, T., Meinl, T., Ohl, P., Thiel, K. and Wiswedel, B., 2009. KNIME-the Konstanz information miner: version 2.0 and beyond. AcM SIGKDD explorations Newsletter, 11(1), pp.26–31.

40. Becht, E., McInnes, L., Healy, J., Dutertre, C.A., Kwok, I.W., Ng, L.G., Ginhoux, F. and Newell, E.W., 2019. Dimensionality reduction for visualizing single-cell data using UMAP. Nature biotechnology, 37(1), pp.38–44.

41. Faure, L., Soldatov, R., Kharchenko, P.V. and Adameyko, I., 2023. scFates: a scalable python package for advanced pseudotime and bifurcation analysis from single-cell data. Bioinformatics, 39(1), p.btac746.

42. Kursa, M.B., Jankowski, A. and Rudnicki, W.R., 2010. Boruta–a system for feature selection. Fundamenta informaticae, 101(4), pp.271–285.

43. Thul, P.J., Åkesson, L., Wiking, M., Mahdessian, D., Geladaki, A., Ait Blal, H., Alm, T., Asplund, A., Björk, L., Breckels, L.M. and Bäckström, A., 2017. A subcellular map of the human proteome. Science, 356(6340), p.eaal3321.

44. Edlind, M.P. and Hsieh, A.C., 2014. PI3K-AKT-mTOR signaling in prostate cancer progression and androgen deprivation therapy resistance. Asian journal of andrology, 16(3), pp.378–386.

45. Castoria, G., Giovannelli, P., Di Donato, M., Hayashi, R., Arra, C., Appella, E., Auricchio, F. and Migliaccio, A., 2013. Targeting androgen receptor/Src complex impairs the aggressive phenotype of human fibrosarcoma cells. PloS one, 8(10), p.e76899.

46. Cleutjens, K.B., van Eekelen, C.C., van der Korput, H.A., Brinkmann, A.O. and Trapman, J., 1996. Two Androgen Response Regions Cooperate in Steroid Hormone Regulated Activity of the Prostate-specific Antigen Promoter (∗). Journal of Biological Chemistry, 271(11), pp.6379–6388.

47. King, C.J., Woodward, J., Schwartzman, J., Coleman, D.J., Lisac, R., Wang, N.J., Van Hook, K., Gao, L., Urrutia, J., Dane, M.A. and Heiser, L.M., 2017. Integrative molecular network analysis identifies emergent enzalutamide resistance mechanisms in prostate cancer. Oncotarget, 8(67), p.111084.

48. Deng, S., Wang, C., Wang, Y., Xu, Y., Li, X., Johnson, N.A., Mukherji, A., Lo, U.G., Xu, L., Gonzalez, J. and Metang, L.A., Ectopic JAK-STAT activation enables the transition to a stem-like and multilineage state conferring AR-targeted therapy resistance. Nat Cancer. 2022; 3 (9): 1071–1087 [online].

49. Xu, P., Yang, J.C., Chen, B., Nip, C., Van Dyke, J.E., Zhang, X., Chen, H.W., Evans, C.P., Murphy, W.J. and Liu, C., 2023. Androgen receptor blockade resistance with enzalutamide in prostate cancer results in immunosuppressive alterations in the tumor immune microenvironment. Journal for Immunotherapy of Cancer, 11(5), p.e006581.

50. Shorning, B.Y., Dass, M.S., Smalley, M.J. and Pearson, H.B., 2020. The PI3K-AKT-mTOR pathway and prostate cancer: at the crossroads of AR, MAPK, and WNT signaling. International journal of molecular sciences, 21(12), p.4507.

51. Claessens, F., Helsen, C., Prekovic, S., Van den Broeck, T., Spans, L., Van Poppel, H. and Joniau, S., 2014. Emerging mechanisms of enzalutamide resistance in prostate cancer. Nature reviews urology, 11(12), pp.712–716.

52. Cham, J., Venkateswaran, A.R. and Bhangoo, M., 2021. Targeting the PI3K-AKT-mTOR pathway in castration resistant prostate cancer: a review article. Clinical Genitourinary Cancer, 19(6), pp.563–e1.

53. Carrion-Salip, D., Panosa, C., Menendez, J.A., Puig, T., Oliveras, G., Pandiella, A., De Llorens, R. and Massaguer, A., 2012. Androgen-independent prostate cancer cells circumvent EGFR inhibition by overexpression of alternative HER receptors and ligands. International journal of oncology, 41(3), pp.1128–1138.

54. Luo, Y., Yang, X., Basourakos, S.P., Zuo, X., Wei, D., Zhao, J., Li, M., Li, Q., Feng, T., Guo, P. and Jiang, Y., 2021. Enzalutamide-resistant progression of castrationresistant prostate cancer is driven via the JAK2/STAT1-dependent pathway. Frontiers in molecular biosciences, 8, p.652443.

55. Chen, W.S., Aggarwal, R., Zhang, L., Zhao, S.G., Thomas, G.V., Beer, T.M., Quigley, D.A., Foye, A., Playdle, D., Huang, J. and Lloyd, P., 2019. Genomic drivers of poor prognosis and enzalutamide resistance in metastatic castration-resistant prostate cancer. European urology, 76(5), pp.562–571.

56. Geng, C., Zhang, M.C., Manyam, G.C., Vykoukal, J.V., Fahrmann, J.F., Peng, S., Wu, C., Park, S., Kondraganti, S., Wang, D. and Robinson, B.D., 2023. SPOP mutations target STING1 signaling in prostate cancer and create therapeutic vulnerabilities to PARP inhibitor–induced growth suppression. Clinical Cancer Research, 29(21), pp.4464–4478.

57. Pearson, H.B., Li, J., Meniel, V.S., Fennell, C.M., Waring, P., Montgomery, K.G., Rebello, R.J., Macpherson, A.A., Koushyar, S., Furic, L. and Cullinane, C., 2018. Identification of Pik3ca mutation as a genetic driver of prostate cancer that cooperates with Pten loss to accelerate progression and castration-resistant growth. Cancer discovery, 8(6), pp.764–779.

58. Vousden, K.H. and Lane, D.P., 2007. p53 in health and disease. Nature reviews Molecular cell biology, 8(4), pp.275–283.

59. Shiau, J.P., Chuang, Y.T., Tang, J.Y., Yang, K.H., Chang, F.R., Hou, M.F., Yen, C.Y. and Chang, H.W., The impact of oxidative stress and AKT pathway on cancer cell functions and its application to natural products. Antioxidants 2022; 11: 1845 [online]

60. Pérez-Sala, D., Oeste, C.L., Martínez, A.E., Carrasco, M.J., Garzón, B. and Canada, F.J., 2015. Vimentin filament organization and stress sensing depend on its single cysteine residue and zinc binding. Nature communications, 6(1), p.7287.

61. Toren, P., Kim, S., Cordonnier, T., Crafter, C., Davies, B.R., Fazli, L., Gleave, M.E. and Zoubeidi, A., 2015. Combination AZD5363 with enzalutamide significantly delays enzalutamide-resistant prostate cancer in preclinical models. European urology, 67(6), pp.986–990.

62. Coleman, D.J., Van Hook, K., King, C.J., Schwartzman, J., Lisac, R., Urrutia, J., Sehrawat, A., Woodward, J., Wang, N.J., Gulati, R. and Thomas, G.V., 2016. Cellular androgen content influences enzalutamide agonism of F877L mutant androgen receptor. Oncotarget, 7(26), p.40690.

63. Nadiminty, N., Tummala, R., Liu, C., Yang, J., Lou, W., Evans, C.P. and Gao, A.C., 2013. NF-κB2/p52 induces resistance to enzalutamide in prostate cancer: role of androgen receptor and its variants. Molecular cancer therapeutics, 12(8), pp.1629–1637.

64. Panja, S., Truica, M.I., Yu, C.Y., Saggurthi, V., Craige, M.W., Whitehead, K., Tuiche, M.V., Al-Saadi, A., Vyas, R., Ganesan, S. and Gohel, S., 2024. Mechanism-centric regulatory network identifies NME2 and MYC programs as markers of Enzalutamide resistance in CRPC. Nature communications, 15(1), p.352.

65. A. Bishop, J.L., Sio, A., Angeles, A., Roberts, M.E., Azad, A.A., Chi, K.N. and Zoubeidi, A., 2014. PD-L1 is highly expressed in Enzalutamide resistant prostate cancer. Oncotarget, 6(1), p.234.

65. Monga, J., Adrianto, I., Rogers, C., Gadgeel, S., Chitale, D., Alumkal, J.J., Beltran, H., Zoubeidi, A. and Ghosh, J., 2022. Tribbles 2 pseudokinase confers enzalutamide resistance in prostate cancer by promoting lineage plasticity. Journal of Biological Chemistry, 298(2), p.101556.

66. Mikolajczyk, S.D., Millar, L.S., Tsinberg, P., Coutts, S.M., Zomorrodi, M., Pham, T., Bischoff, F.Z. and Pircher, T.J., 2011. Detection of EpCAM-negative and cytokeratin-negative circulating tumor cells in peripheral blood. Journal of oncology, 2011(1), p.252361.

67. Lazar, D.C., Cho, E.H., Luttgen, M.S., Metzner, T.J., Uson, M.L., Torrey, M., Gross, M.E. and Kuhn, P., 2012. Cytometric comparisons between circulating tumor cells from prostate cancer patients and the prostate-tumor-derived LNCaP cell line. Physical biology, 9(1), p.016002.

68. Hyun, K.A., Koo, G.B., Han, H., Sohn, J., Choi, W., Kim, S.I., Jung, H.I. and Kim, Y.S., 2016. Epithelial-to-mesenchymal transition leads to loss of EpCAM and different physical properties in circulating tumor cells from metastatic breast cancer. Oncotarget, 7(17), p.24677.

69. Fallahi-Sichani, M., Honarnejad, S., Heiser, L.M., Gray, J.W. and Sorger, P.K., 2013. Metrics other than potency reveal systematic variation in responses to cancer drugs. Nature chemical biology, 9(11), pp.708–714.

70. Thompson-Elliott, B., Johnson, R. and Khan, S.A., 2021. Alterations in TGFβ signaling during prostate cancer progression. American journal of clinical and experimental urology, 9(4), p.318.

71. Huang, G., Osmulski, P.A., Bouamar, H., Mahalingam, D., Lin, C.L., Liss, M.A., Kumar, A.P., Chen, C.L., Thompson, I.M., Sun, L.Z. and Gaczynska, M.E., 2016. TGF-β signal rewiring sustains epithelial-mesenchymal transition of circulating tumor cells in prostate cancer xenograft hosts. Oncotarget, 7(47), p.77124.

72. Small, E.J., Huang, J., Youngren, J., Sokolov, A., Aggarwal, R.R., Thomas, G., True, L.D., Zhang, L., Foye, A., Alumkal, J.J. and Ryan, C.J., 2015. Characterization of neuroendocrine prostate cancer (NEPC) in patients with metastatic castration resistant prostate cancer (mCRPC) resistant to abiraterone (Abi) or enzalutamide (Enz): Preliminary results from the SU2C/PCF/AACR West Coast Prostate Cancer Dream Team (WCDT).

73. Luo, M., Huang, Z., Yang, X., Chen, Y., Jiang, J., Zhang, L., Zhou, L., Qin, S., Jin, P., Fu, S. and Peng, L., 2022. PHLDB2 mediates cetuximab resistance via interacting with EGFR in latent metastasis of colorectal cancer. Cellular and Molecular Gastroenterology and Hepatology, 13(4), pp.1223–1242.

74. Xie, Y., Ning, S. and Hu, J., 2022. Molecular mechanisms of neuroendocrine differentiation in prostate cancer progression. Journal of Cancer Research and Clinical Oncology, 148(7), pp.1813–1823.

75. Edwards, J., Krishna, N.S., Grigor, K.M. and Bartlett, J.M.S., 2003. Androgen receptor gene amplification and protein expression in hormone refractory prostate cancer. British journal of cancer, 89(3), pp.552–556.

76. Bonomi, P.D., Gandara, D., Hirsch, F.R., Kerr, K.M., Obasaju, C., Paz-Ares, L., Bellomo, C., Bradley, J.D., Bunn Jr, P.A., Culligan, M. and Jett, J.R., 2018. Predictive biomarkers for response to EGFR-directed monoclonal antibodies for advanced squamous cell lung cancer. Annals of Oncology, 29(8), pp.1701–1709.

77. Ahn, S., Woo, J.W., Lee, K. and Park, S.Y., 2020. HER2 status in breast cancer: changes in guidelines and complicating factors for interpretation. Journal of pathology and translational medicine, 54(1), pp.34–44.

78. Chen, X.J., Yuan, S.Q., Duan, J.L., Chen, Y.M., Chen, S., Wang, Y. and Li, Y.F., 2020. The value of PD-L1 expression in predicting the efficacy of anti-PD-1 or anti-PD-L1 therapy in patients with cancer: a systematic review and meta-analysis. Disease Markers, 2020(1), p.6717912.

79. Seifert, R., Seitzer, K., Herrmann, K., Kessel, K., Schäfers, M., Kleesiek, J., Weckesser, M., Boegemann, M. and Rahbar, K., 2020. Analysis of PSMA expression and outcome in patients with advanced prostate cancer receiving 177Lu-PSMA-617 radioligand therapy. Theranostics, 10(17), p.7812.

80. Janku, F., Wheler, J.J., Naing, A., Falchook, G.S., Hong, D.S., Stepanek, V.M., Fu, S., Piha-Paul, S.A., Lee, J.J., Luthra, R. and Tsimberidou, A.M., 2013. PIK3CA mutation H1047R is associated with response to PI3K/AKT/mTOR signaling pathway inhibitors in early-phase clinical trials. Cancer research, 73(1), pp.276–284.

81. Zhang, X., Park, J.S., Park, K.H., Kim, K.H., Jung, M., Chung, H.C., Rha, S.Y. and Kim, H.S., 2015. PTEN deficiency as a predictive biomarker of resistance to HER2-targeted therapy in advanced gastric cancer. Oncology, 88(2), pp.76–85.

82. Fasching, P.A., Heusinger, K., Haeberle, L., Niklos, M., Hein, A., Bayer, C.M., Rauh, C., Schulz-Wendtland, R., Bani, M.R., Schrauder, M. and Kahmann, L., 2011. Ki67, chemotherapy response, and prognosis in breast cancer patients receiving neoadjuvant treatment. BMC cancer, 11, pp.1–13.

83. Turnham, D.J., Bullock, N., Dass, M.S., Staffurth, J.N. and Pearson, H.B., 2020. The PTEN conundrum: how to target PTEN-deficient prostate cancer. Cells 9, 2342 [online].

84. Shin, S.H., Bode, A.M. & Dong, Z. Precision medicine: the foundation of future cancer therapeutics. npj Precision Onc 1, 12 (2017). 10.1038/s41698-017-0016-z

85. Lassen, U. N., Makaroff, L. E., Stenzinger, A., Italiano, A., Vassal, G., Garcia-Foncillas, J., & Avouac, B. (2021). Precision Oncology: a Clinical and Patient Perspective. Future Oncology, 17(30), 3995–4009. 10.2217/fon-2021-0688

86. Ku, SY., Gleave, M.E. & Beltran, H. Towards precision oncology in advanced prostate cancer. Nat Rev Urol 16, 645–654 (2019). 10.1038/s41585-019-0237-8

87. Keller, L., Pantel, K. Unravelling tumour heterogeneity by single-cell profiling of circulating tumour cells. Nat Rev Cancer 19, 553–567 (2019).

88. Tsao, S.C.H., Wang, J., Wang, Y., Behren, A., Cebon, J. and Trau, M., 2018. Characterising the phenotypic evolution of circulating tumour cells during treatment. Nature communications, 9(1), p.1482.

89. Alix-Panabières, C., Mader, S. and Pantel, K., 2017. Epithelial-mesenchymal plasticity in circulating tumor cells. Journal of Molecular Medicine, 95(2), pp.133–142.

90. Ma, Y., Luk, A., Young, F.P., Lynch, D., Chua, W., Balakrishnar, B., De Souza, P. and Becker, T.M., 2016. Droplet digital PCR based androgen receptor variant 7 (AR-V7) detection from prostate cancer patient blood biopsies. International journal of molecular sciences, 17(8), p.1264.

91. Quintanal-Villalonga, Á., Chan, J.M., Yu, H.A., Pe’er, D., Sawyers, C.L., Sen, T. and Rudin, C.M., 2020. Lineage plasticity in cancer: a shared pathway of therapeutic resistance. Nature reviews Clinical oncology, 17(6), pp.360–371.

92. Paschalis, A., Sheehan, B., Riisnaes, R., Rodrigues, D.N., Gurel, B., Bertan, C., Ferreira, A., Lambros, M.B., Seed, G., Yuan, W. and Dolling, D., 2019. Prostate-specific membrane antigen heterogeneity and DNA repair defects in prostate cancer. European urology, 76(4), pp.469–478.

93. McGranahan, N. and Swanton, C., 2017. Clonal heterogeneity and tumor evolution: past, present, and the future. Cell, 168(4), pp.613–628.

94. Zhang, C., Guan, Y., Sun, Y., Ai, D. and Guo, Q., 2016. Tumor heterogeneity and circulating tumor cells. Cancer letters, 374(2), pp.216–223.

95. Tellez-Gabriel, M., Heymann, M.F. and Heymann, D., 2019. Circulating tumor cells as a tool for assessing tumor heterogeneity. Theranostics, 9(16), p.4580.

96. Tang, C., Fu, S., Jin, X., Li, W., Xing, F., Duan, B., Cheng, X., Chen, X., Wang, S., Zhu, C. and Li, G., 2023. Personalized tumor combination therapy optimization using the single-cell transcriptome. Genome Medicine, 15(1), p.105.

97. Plana, D., Palmer, A.C. and Sorger, P.K., 2022. Independent drug action in combination therapy: implications for precision oncology. Cancer discovery, 12(3), pp.606–624.

98. Hein, M.Y., Peng, D., Todorova, V., McCarthy, F., Kim, K., Liu, C., Savy, L., Januel, C., Baltazar-Nunez, R., Sekhar, M. and Vaid, S., 2025. Global organelle profiling reveals subcellular localization and remodeling at proteome scale. Cell, 188(4), pp.1137–1155.

99. Xue, Z.Z., Wu, Y., Gao, Q.Z., Zhao, L. and Xu, Y.Y., 2020. Automated classification of protein subcellular localization in immunohistochemistry images to reveal biomarkers in colon cancer. BMC bioinformatics, 21(1), p.398.

